# A comprehensive molecular atlas of the cell types in the mouse liver

**DOI:** 10.1101/2025.01.24.634721

**Authors:** Riikka Pietilä, Guillem Genové, Giuseppe Mocci, Yuyang Miao, Jianping Liu, Stefanos Leptidis, Francesca Del Gaudio, Martin Uhrbom, Elisa Vázquez-Liébanas, Sonja Gustafsson, Byambajav Buyandelger, Elisabeth Raschperger, Johan LM. Björkegren, Emil M. Hansson, Konstantin Gaengel, Maarja Andaloussi Mäe, Marie Jeansson, Michael Vanlandewijck, Liqun He, Carina Strell, Xiao-Rong Peng, Urban Lendahl, Christer Betsholtz, Lars Muhl

## Abstract

The liver plays a crucial role in essential physiological processes, and its impaired function due to liver fibrosis from various causes is an increasingly significant health issue. The liver’s functionality relies on the precise arrangement of its cellular structures, yet the molecular architecture of these units remains only partially understood. We created a comprehensive molecular atlas detailing all the major cell types present in the adult mouse liver through deep single-cell RNA sequencing. Our analysis offers new insights into hepatic endothelial and mesenchymal cells, specifically highlighting the differences between the cells of the periportal microvasculature, the sinusoids, and the portal vein, the latter exhibiting a mixed arterio-venous phenotype. We identified distinct subpopulations of hepatic stellate cells, fibroblasts, and vascular mural cells located in different anatomical regions. Comparisons with transcriptomic data from disease models indicate that a previously unrecognized capsular population of hepatic stellate cells expands in response to fibrotic disease. Our findings reveal that various fibroblast subpopulations respond differently to pathological insults. This data resource will be invaluable for advancing therapeutic interventions targeting hepatic diseases.

## Introduction

The liver performs many critical physiological functions, including metabolic regulation, immune surveillance, and the detoxification of harmful metabolites and toxins (Shetty *et al*, 2018). Structurally, the liver is comprised of lobules, each featuring a specific arrangement of cells that distribute and function along an axis extending from the peripheral portal area (portal triad) to the central vein (Figure 1A). Hepatocytes, the primary parenchymal cells of the liver, are aligned along this portal-central axis and carry out various metabolic and synthetic functions, including the production of plasma proteins and bile (Halpern *et al*, 2017).

**Figure 1:**
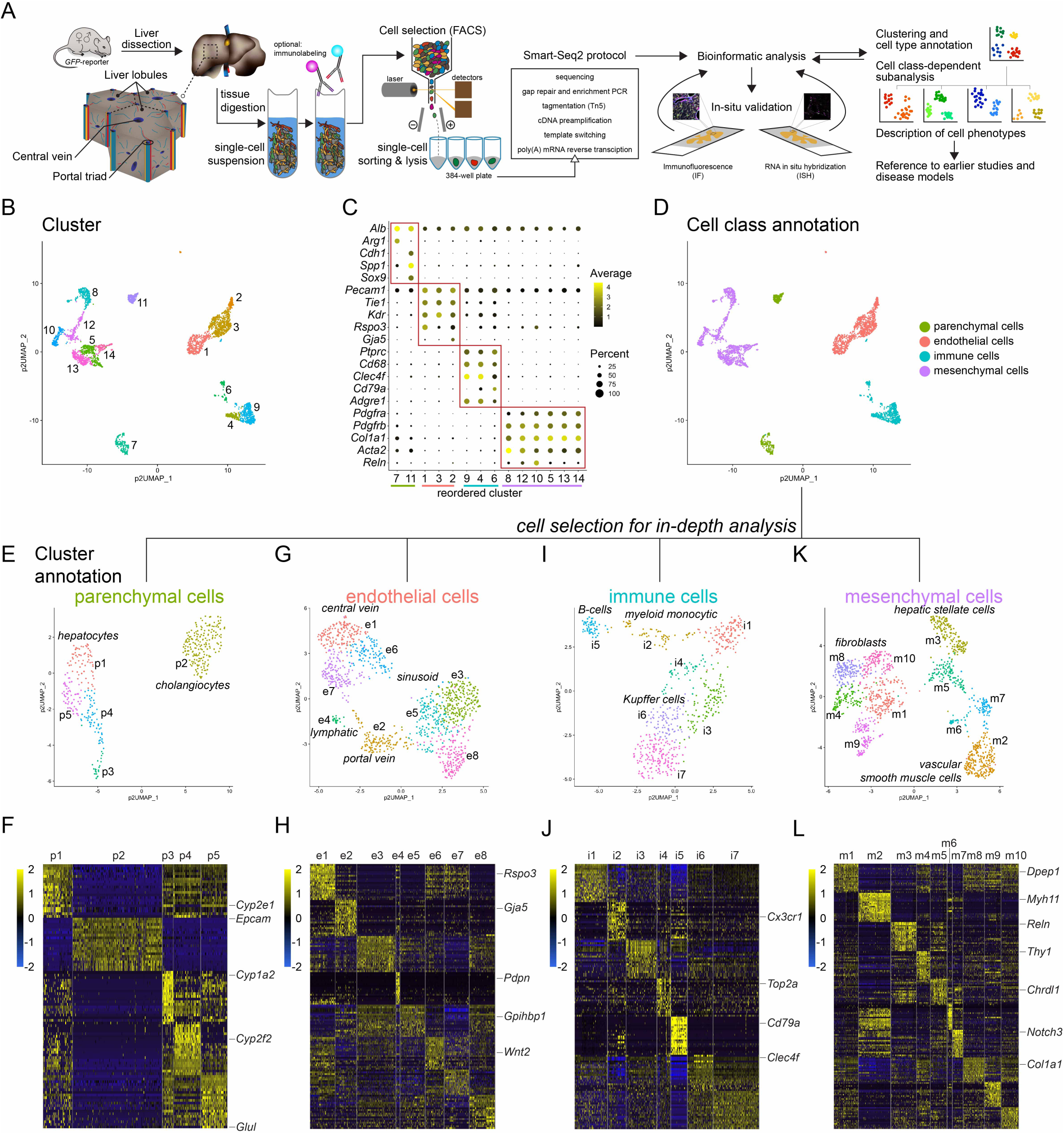
Liver cells characterized by scRNA-seq. **A** Schematic overview of the experimental layout. **B** UMAP visualization of the clustering result using pagoda2 multilevel setting of the complete adult mouse liver scRNA-seq dataset (in total 3491 single-cell transcriptomes), annotated and color coded for the different clusters. **C** Dot plot showing the expression of canonical marker genes to define cell type classes. **D** UMAP visualization of the complete dataset color coded for cell type classes: parenchymal cells, endothelial cells, immune cells, and mesenchymal cells. **E-L** UMAP visualization of the clustering results and heat maps showing the top20 cluster-enriched genes in the separate analysis of the parenchymal cell (E,F), endothelial cell (G,H), immune cell (I,J), and mesenchymal cell (K,L) datasets.

The vascular system of the liver displays unique characteristics, notably a dual blood supply comprising (i) nutrient-rich, oxygen-poor blood from the portal vein, which drains the gut, and (ii) oxygen-rich blood from the hepatic artery, a branch from the celiac trunk of the aorta. These two blood supplies converge in the hepatic capillary system, known as the sinusoids, which drain into the central vein. The sinusoids are highly specialized and distinguish in multiple ways from capillaries in most other organs. Their distinct features include discontinuous endothelial lining, which allows macromolecules to exchange freely between the blood and parenchyma (Sorensen *et al*, 2015), as well as the lack of a distinct basement membrane and associated regular pericytes.

The space between the sinusoidal endothelial wall and the hepatocytes harbors two liver-specific cell types, the hepatic stellate cells (HSC), which are mesenchymal cells with properties similar to both pericytes and fibroblasts (Kamm & McCommis, 2022), and the Kupffer cells, which are a type of resident macrophage (Guilliams & Scott, 2022). Additionally, the liver contains the cholangiocyte-lined bile duct system (Tabibian *et al*, 2013), which runs parallel to the portal veins, venules, and hepatic arteries. Fibroblasts in the connective tissue surrounding the portal triad, together with mesothelial cells at the liver capsule, contribute to the formation of hepatic fibrous tissue sheaths also known as *the tunics of Glisson* (Helling & McCleary, 2016; Wells, 2014a).

While the anatomy and histology of the liver is well-established, in-depth information about how gene expression patterns differ among the various specialized hepatic cell types under physiological conditions and in disease has only recently been unraveled, primarily due to the advent of single-cell RNA-sequencing (scRNA-seq) and spatial transcriptomic techniques (Dobie *et al*, 2019; Halpern *et al*, 2018; Halpern *et al*., 2017; Hildebrandt *et al*, 2021; MacParland *et al*, 2018; Tabula Muris *et al*, 2018; Watson *et al*, 2025). Despite this progress, detailed transcriptomic information covering all major hepatic cell types at homeostasis remains limited, but would be valuable as baseline for comparison to transcriptomic data from pathological conditions. It would be important to establish the precise molecular profiles for several less noticed hepatic cell types, including vascular mural cells, fibroblasts, and cholangiocytes. For example, it remains largely unexplored how HSC compare to their closest relatives in other organs; the vascular mural cells. Moreover, a parallel analysis of the different mesenchymal cell populations of the liver has not been conducted. Such information will be crucial for fully understanding the liver’s cellular architecture and how hepatic cells respond to injury and disease. A more complete inventory of hepatic cell types will also illuminate potential hepatic transcriptional zonation (gene expression differences in the same cell type along an anatomical axis) for cell types other than hepatocytes and sinusoidal endothelial cells, for which zonation has been described along the portal-central axis, as well as for HSC, for which zonation has been suggested (Dobie *et al*., 2019; Duan *et al*, 2022; Guilliams & Scott, 2022; Halpern *et al*., 2018; Halpern *et al*., 2017; Inverso *et al*, 2021; Krenkel *et al*, 2019; Paris & Henderson, 2022; Rosenthal *et al*, 2021). The gradual or punctuated phenotypic variation of other mesenchymal hepatic cell types along spatial axes remains less explored.

To address these questions, we have generated a comprehensive molecular atlas of the various parenchymal and mesenchymal cell types of the adult mouse liver, utilizing a deep scRNA-seq approach. This, combined with immunofluorescence (IF) and *in situ* RNA hybridization (ISH), helped establishing genome-wide transcriptomic profiles for all major hepatic cell types. It also identified several hitherto poorly characterized cell type subpopulations in the liver elucidating the zonation principles for HSC, vascular smooth muscle (mural) cells, and fibroblasts. With this transcriptomic and anatomical information at hand, we reevaluated data from several previously published hepatic scRNA-seq studies, encompassing normal and diseased livers from both mice and humans, to formulate refined conclusions about disease-regulated gene expression patterns in distinct mesenchymal cell subpopulations.

## Results

### Transcriptional profiling of the principal cell types of the adult mouse liver

To obtain single-cell transcriptomic information from all major cell types in the adult mouse liver, we employed a combination of transgenic reporter mice and antibody-based cell enrichment strategies, as schematically depicted in Figure 1A, and detailed in the Methods section. Cholangiocytes were isolated using antibodies against EPCAM (epithelial cell adhesion molecule, CD326), while hepatocytes could be captured in sufficient numbers without enrichment, due to their abundance in liver tissue. Endothelial cells were isolated by immunopanning with antibodies against PECAM1 (platelet/endothelial cell adhesion molecule 1, CD31) and/or VE-cadherin (cadherin 5, CDH5/CD144). Kupffer cells were collected using an antibody against CD68. Transgenic reporter mice expressing GFP from the promoter constructs from *Pdgfrb* (*Pdgfrb^GFP^*), *Pdgfra* (*Pdgfra^H2bGFP^*), or *Acta2* (*Acta2^GFP^*) were utilized to enrich for different mesenchymal cell populations, as previously described (Muhl *et al*, 2020; Muhl *et al*, 2022b). In total, we analyzed the transcriptomes of 3,491 cells derived from livers of adult C57Bl6 mice of both sexes. The Smart-seq2 protocol was employed to achieve the deepest possible mRNA sequence capture from each individual cell (Picelli *et al*, 2014) (Figure 1A).

A first round of combined analysis of all captured cells using the Seurat analysis pipeline (Satija *et al*, 2015; Stuart *et al*, 2019), combined with cluster definitions based on the pagoda2 algorithm (Fan *et al*, 2016), revealed 14 distinct clusters (Figure 1B). By assessing the expression of widely accepted canonical marker genes for specific cell types and classes (Figure 1C, Figure S1A), we grouped these 14 clusters into four distinct cell classes encompassing parenchymal cells, endothelial cells, immune cells, and mesenchymal cells (Figure 1D). To obtain a more granular view of each cell class, we separately analyzed and further annotated them (Figure 1E-L). The parenchymal cells split into five distinguishable clusters comprising one cluster of cholangiocytes and four clusters of hepatocytes (Figure 1E,F). Endothelial cells separated into eight clusters, comprising subtypes originating from the portal vein, capillary sinusoids, central vein and lymphatics (Figure 1G,H). Immune cells distributed into seven clusters, representing B-cells, myeloid monocytic cells, and Kupffer cells/macrophages (Figure 1I,J). Mesenchymal cells formed ten clusters, encompassing different types of fibroblasts, HSC, and vascular smooth muscle cells (SMC) (Figure 1K,L).

From the detailed analysis, as outlined in the following chapters, we constructed an accompanying, searchable database including UMAP visualization of the complete and the four separately analyzed datasets, along with bar plot-visualization of the complete dataset (Figure S1B,C), This database can be explored cell-by-cell and gene-by-gene at https://betsholtzlab.org/Publications/LiverScRNAseq/search.html.

### Parenchymal cells

Hepatocytes are organized as stacks of repetitive cellular layers within the liver lobules. Within each layer, hepatocyte zonation along the portal-central axis has been reported based on morphological observations (Asada-Kubota & Kanamura, 1981; Rappaport & Potvin, 1963) and transcriptomic analysis of single-cells (Aizarani *et al*, 2019; Halpern *et al*., 2017). Our data identified four hepatocyte clusters (cluster #p1, p3, p4, p5) (Figure S2A), corroborating earlier reports of a transcriptomic hepatocyte zonation along the portal-central axis, but further revealed a distinct hepatocyte population that expressed *Lgr5* and located pericentrally (Planas-Paz *et al*, 2016) (Figure S2A-D).

Cholangiocytes constitute the second major type of parenchymal cells in the liver (Banales *et al*, 2019). Somewhat unexpectedly, we identified only one cholangiocyte cluster (cluster #p2) (Figure S2A), despite the previously reported heterogeneity in cholangiocyte cell size (Fukushima & Ueno, 2006; Li *et al*, 2023; Tabibian *et al*., 2013; Tulasi *et al*, 2021; Ueno *et al*, 2003). Cholangiocyte-enriched transcripts encoded for example multiple claudins, cytokeratins, and solute carrier transporters (Figure S2E,F).

Our present analysis largely confirms the previously described heterogeneity and zonation of the hepatic parenchyma (Halpern *et al*., 2017). We therefore chose to focus our analysis on hepatic stromal cells, as detailed in the following chapters.

### Endothelial cells

In parallel to the zonation observed in hepatocytes, sinusoidal endothelial cells also exhibit a zonation pattern (Duan *et al*., 2022; Halpern *et al*., 2018; Inverso *et al*., 2021). Separate analysis of the 1,106 endothelial cell transcriptomes identified eight distinguishable clusters that confirm the previously reported molecular zonation along the portal-central axis. Annotation based on commonly accepted markers (Figure 1H) assigned one cluster (cluster #e2) as portal vein endothelial cells, four clusters (clusters #e3, e5, e6 and e8) as sinusoidal capillary endothelial cells, and two clusters (clusters #e1 and e7) as central vein endothelial cells (Figure 2A,B). The cluster of portal vein endothelial cells could be split further into two clusters (#e2c and e5c), using the pagoda2 *community* setting (Figure 2A,C, Figure S2G). Endothelial cell cluster #e4/e4c represents lymphatic endothelial cells identified by the expression of canonical markers (Ulvmar & Makinen, 2016), such as *Prox1*, *Mmrn1*, *Pdpn*, *Thy1*, and *Ccl21a* (Figure S2H); these cells will not be further discussed herein.

**Figure 2:**
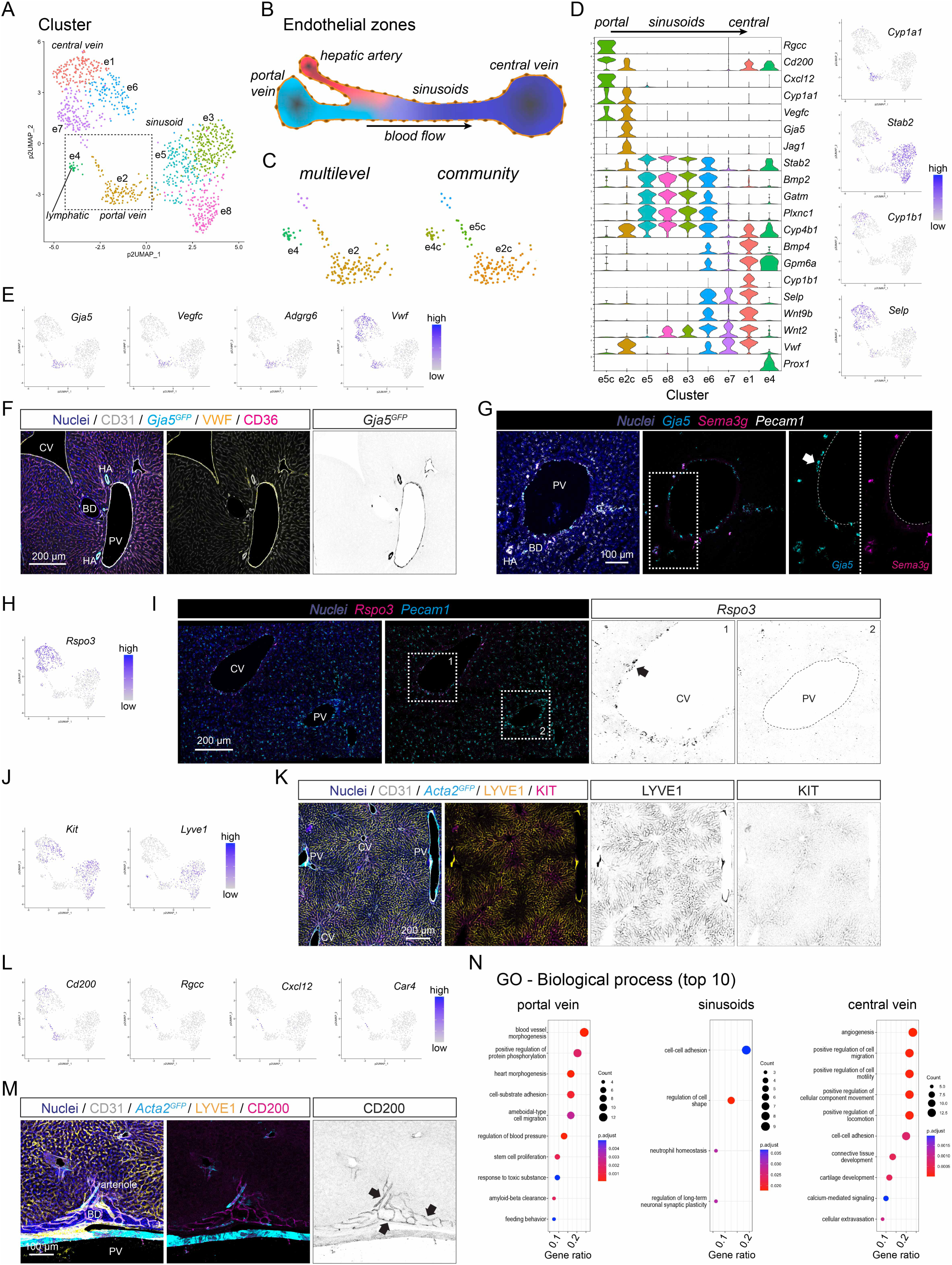
Endothelial cell subset analysis. **A** UMAP visualization of pagoda2 clustering result of the endothelial cell dataset. **B** Schematic depiction of the intra-hepatic vasculature. **C** Magnified section of the UMAP landscape (indicated in A) showing the pagoda2 clustering result using the multilevel (left panel) or community (right panel) setting, respectively. **D** Violin plot showing genes with differential expression in endothelial cells along the portal-central axis. **E** UMAP visualization of the expression levels of *Gja5*, *Vegfc*, *Adgrg6*, and *Vwf*. **F** IF for CD31, VWF, and CD36 on a liver tissue section from a *Gja5^GFP^* reporter mouse. **G** ISH for *Gja5*, *Sema3g*, and *Pecam1* on a liver tissue section. **H** UMAP visualization of the expression levels of *Rspo3*. **I** ISH for *Rspo3* and *Pecam1* on a liver tissue section. **J** UMAP visualization of the expression levels of *Kit* and *Lyve1*. **K** IF for CD31, LYVE1, and KIT on a liver tissue section from an *Acta2^GFP^* reporter mouse. **L** UMAP visualization of the expression levels of *Cd200, Cxcl12*, *Rgcc*, and *Car4*. **M** IF for CD200, CD31, and LYVE1 on a liver tissue section from an *Acta2^GFP^* reporter mouse. **N** Dot plot showing the top10 overrepresented GO terms for the endothelial cell subsets (portal vein, left panel; sinusoids, middle panel; central vein, right panel). PV: portal vein, CV: central vein, HA: hepatic artery, BD: bile duct. Scale bars are indicated in the respective image panels.

Differential gene expression (DEG) analysis revealed zone-specific gene expression patterns. For example, transcripts for *Vegfc*, *Cyp1a1* (encoding a cytochrome P450 family member monooxygenase, also known as aryl hydrocarbon hydroxylase, AHH), *Gja5*, and *Jag1* were enriched in portal vein endothelial cells, *Stab2*, *Bmp2*, *Gatm*, and *Plxnc1* in sinusoidal endothelial cells, and *Bmp4*, *Cyp1b1*, and *Selp* (encoding P-selectin) in central vein endothelial cells (Figure 2D). ISH for *Adgrg6* (encoding the adhesion G protein-coupled receptor G6) and *Cyp1a1* (suggested to protect against metabolic hepatic diseases) (Uno *et al*, 2018), confirmed that cells in cluster #e2c originate from the portal vein, rather than the hepatic artery (Figure S2I,J). We next analyzed the localization of *Gja5* (encoding the gap junction protein connexin 40), a known marker for arterial endothelial cells (Bastide *et al*, 1993) (Figure 2E). Analysis of the *Gja5^GFP^* reporter mice (Miquerol *et al*, 2004) revealed a strong GFP signal in both hepatic artery and portal vein endothelial cells, but not in the sinusoids or central veins (Figure 2F). ISH confirmed the presence of *Gja5* mRNA in portal vein endothelial cells (Figure 2G, Figure S2K). Because *Gja5* is a canonical marker of arterial endothelial cells in many organs, its expression in the portal vein suggests that these cells have a mixed phenotype, including expression of both venous and arterial marker genes. ISH for another arterial endothelial cell marker gene *Sema3g* (Kutschera *et al*, 2011) showed high expression in endothelial cells lining hepatic arteries and peribiliary arterioles (Figure 2G, Figure S2K). However, the transcript for *Sema3g* was not found in our scRNA-seq database, suggesting that hepatic artery endothelial cells may not have been captured for scRNA-seq in our experiments. ISH confirmed the specific expression of *Rspo3* by endothelial cells in and around the central vein, but not at the portal tract (Figure 2H,I). IF analysis for KIT and LYVE1 revealed expression in sinusoidal endothelial cells, with KIT being localized pericentrally and LYVE1 exhibiting mid-sinusoidal expression (Figure 2J,K, Figure S2L). In addition to the zonal endothelial cell gene expression pattern along the portal-central axis, a small cluster, #e5c, identified by the community setting of pagoda2, mapped between the portal and central vein but distant from the sinusoidal endothelial cells in the UMAP display (Figure 2C). Cells in cluster #e5c are characterized by expression of *Cd200*, a suggested regulator of immune and inflammatory processes (Choe & Choi, 2023), *Rgcc* (encoding the regulator of cell cycle), *Cxcl12*, and *Car4* (encoding the carbonic anhydrase 4) previously shown with specific expression in a subset of lung capillary endothelial cells (Gillich *et al*, 2020) (Figure 2L, Figure S2M). IF for CD200 showed expression in capillaries and arterioles of the peribiliary vascular plexus (Figure 2M) (Morell *et al*, 2013), suggesting that the cells in the cluster #e5c likely originate from the peribiliary vasculature (PBV). Transcripts for *Lrg1*, *Ephx1*, and the two elastic fiber constituents *Fbln5* (encoding fibulin 5), and *Eln* (encoding elastin) were enriched in both portal and central vein, in contrast to *Cxcl10* and *Cyp4b1*, which were enriched in sinusoidal endothelium (Figure S2M).

GO analysis of endothelial zone-enriched genes identified terms such as ‘blood-vessel morphogenesis’, ‘regulation of blood pressure’, and ‘response to toxic substance’ associated with portal vein endothelial cells, and terms including ‘cell-cell adhesion’ and ‘neutrophil homeostasis’ with sinusoidal endothelial cells. Terms related to ‘cell migration’, ‘connective tissue development’, and ‘angiogenesis’ were associated with central vein endothelial cells (Figure 2N).

### Immune cells

The liver contains at least two types of immune cells: tissue-resident macrophages (Kupffer cells), and monocyte-derived macrophages (MoMFs) (Guillot & Tacke, 2019; Heymann & Tacke, 2016; Su *et al*, 2021). Kupffer cells constitute the largest population of tissue-resident macrophages in the body and localize across the liver lobule, patrolling/interfacing both with the sinusoidal lumen and the space of Disse (Kubes & Jenne, 2018). Kupffer cells are established in the early embryo and are maintained in adulthood independently of bone-marrow derived monocytes (Yona *et al*, 2013). In contrast, MoMFs, as their name suggests, are macrophages derived from monocytes. They are found in smaller numbers compared to Kupffer cells and primarily localize near the portal tract (English *et al*, 2022; Guilliams & Scott, 2022).

Analysis of a total of 651 transcriptomes from immune cells revealed that most of them (528 cells) represented Kupffer cells (clusters #i1, i3, i4, i6, i7), as determined by the expression of *Clec4f*, *Folr2*, and *Vsig4* (Guilliams *et al*, 2022). Additionally, we identified *Cd79a*-positive B-cells (cluster #i5), as well as a scattered cluster #i2 of myeloid monocytic cells (Figure 3A-C, Figure S3A). Using the pagoda2 community setting, we further subdivided cluster #i2 into three subclusters #i2c, i4c, i7c (Figure 3B, Figure S3B) characterized by the expression of *Siglech* and *Ccr9* markers for dendritic cells (cluster #i7c), *Cd9* and *Cx3cr1* markers for MoMF (cluster #i2c), and *Ddx60* for a subpopulation of Kupffer cell (cluster #i4c) (Figure 3C). Next, we used IF analysis distinguish Kupffer cells from MoMFs (Figure 3D). Cells double positive for CD68 and CLEC4F (Kuppfer cells) were found along the sinusoidal bed, whereas CD68 positive and CLEC4F negative cells (MoMFs) resided near the portal tract (Figure 3E), supporting the previously proposed anatomical distribution of these cell types (English *et al*., 2022). The separation of the Kupffer cells into five subclusters using pagoda2 did however not reveal distinct markers for putative Kupffer cell subtypes, except for a signature associated with cell cycle progression (*Mki67*, *Top2a*, *Ccna2*, *Ccnb1*) in cluster #i4/i5c, and the contamination by endothelial cell transcriptomes (e.g. *Tie1*, *Kdr*, *Ptprb*) in subcluster #i9c (Figure S3A-C). Although earlier studies identified two distinct Kupffer cell populations (designated KC1 & KC2) based on differential expression levels of *Mrc1* (encoding the C-type 1 mannose receptor CD206), *Esam*, and *Cdh5* (the two latter know endothelial cells markers) (Bleriot *et al*, 2021; De Simone *et al*, 2021), all Kupffer cells in our dataset exhibited high expression of *Mrc1* and *Cdh5* while being negative for *Esam*. Because we did not recapitulate the previously described KC1/KC2 subtypes and further noticed that endothelial cell contamination could be a driver of sub-clustering (Figure S3C) (Guilliams & Scott, 2022; Hume *et al*, 2022), we removed the cells corresponding to cluster #i5c (proliferating Kupffer cells) and i9c (Kupffer cells contaminated by endothelial cells) from the consensus Kupffer cell population (containing clusters #i1c, i3c, i8c, i10c, i11c) displayed in the searchable bar plot database (Figure S1B,C).

**Figure 3:**
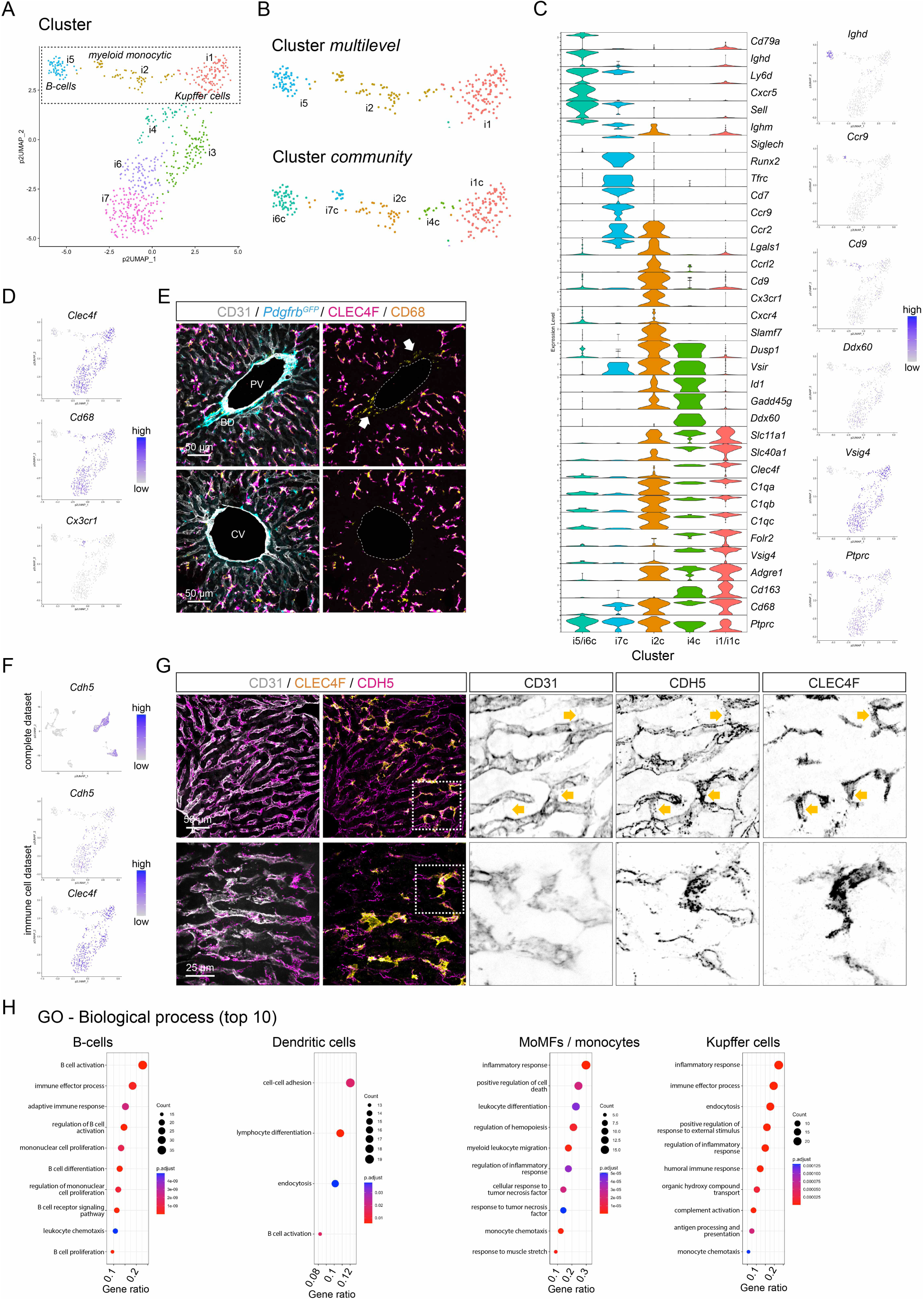
Immune cell subset analysis. **A** UMAP visualization of pagoda2 clustering result of the immune cell dataset. **B** Magnified section of the UMAP landscape (indicated in A) showing the pagoda2 clustering result using the multilevel (upper panel) or community (lower panel) setting, respectively. **C** Violin plot and UMAP visualization showing the expression of enriched genes in the clusters obtained with the community setting. **D** UMAP visualization of the expression levels of *Clec4f*, *Cd68*, and *Cx3cr1*. **E** IF for CD31, CLEC4F, and CD68 on a liver tissue section from a *Pdgfra^H2bGFP^* reporter mouse. The arrows highlight CLEC4F-negative, CD68-positive macrophages. **F** UMAP visualization of the expression level of *Cdh5* in the complete dataset (upper panel), or *Cdh5* and *Clec4f* in the immune cell dataset (lower panel). **G** IF for CD31, CDH5, and CLEC4F on a liver tissue section. The boxed areas are shown magnified to the right. The arrows highlight CDH5 signals overlapping with CLEC4F. **H** Dot plots showing the top10 GO terms overrepresented in genes with enriched expression in (from left to right) B-cells, myeloid monocytic/dendritic cells, MoMFs, or Kupffer cells. PV: portal vein, BD: bile duct, CV: central vein. Scale bars are indicated in the respective image panels.

Notwithstanding the suspected endothelial cell contamination of a small population of the Kupffer cells, our data clearly demonstrated that Kupffer cells express *Cdh5* (encoding cadherin 5, CDH5, also known as vascular endothelial (VE)-cadherin) at levels comparable to those in hepatic endothelial cells (Figure 3F). CDH5 mediates homotypic cell-cell binding, thus suggesting a potential junctional interaction through CDH5 between Kupffer cells and sinusoidal endothelial cells (Ito *et al*, 1980). High-resolution confocal imaging revealed a complex pattern for CDH5 protein localization in sinusoidal endothelial cells depending on their vicinity to Kupffer cells (Figure 3G, Figure S3D).

GO analysis of population-enriched genes confirmed the identities of B-cells and dendritic cells. Additionally, this analysis revealed that the terms ‘inflammatory response’ and ‘monocyte chemotaxis’ were associated with both MoMF and Kupffer cell-enriched genes, suggesting the expression of distinct inflammatory gene expression programs in the two cell types (Figure 3H, Figure S3E).

### Mesenchymal cells

The liver is known to contain multiple mesenchymal cell populations at distinct anatomical locations. HSC, which partially resemble pericytes found in other organs, populate the perisinusoidal space. Vascular SMC and other vascular mural cells reside in the vessel wall of hepatic arteries, arterioles, portal veins, and central veins. Fibroblasts reside in the portal tract and at the liver capsule (Bhunchet & Wake, 1992; Wells, 2014a). In our mesenchymal cell dataset, transcriptomes from in total 1,337 cells from adult mice were classified into ten clusters using the pagoda2 multilevel algorithm (Figure 4A, Figure S4A). HSC, vascular mural cells, and fibroblasts, identified by canonical markers (Figure 1L), were allocated to separate cell clusters within the UMAP display. The distribution suggested heterogeneity not only between, but also within, these cell classes, particularly among the fibroblasts and mural cells (Figure 4A). In the following sections, we present the mesenchymal cell annotations in relation to their anatomical locations.

**Figure 4:**
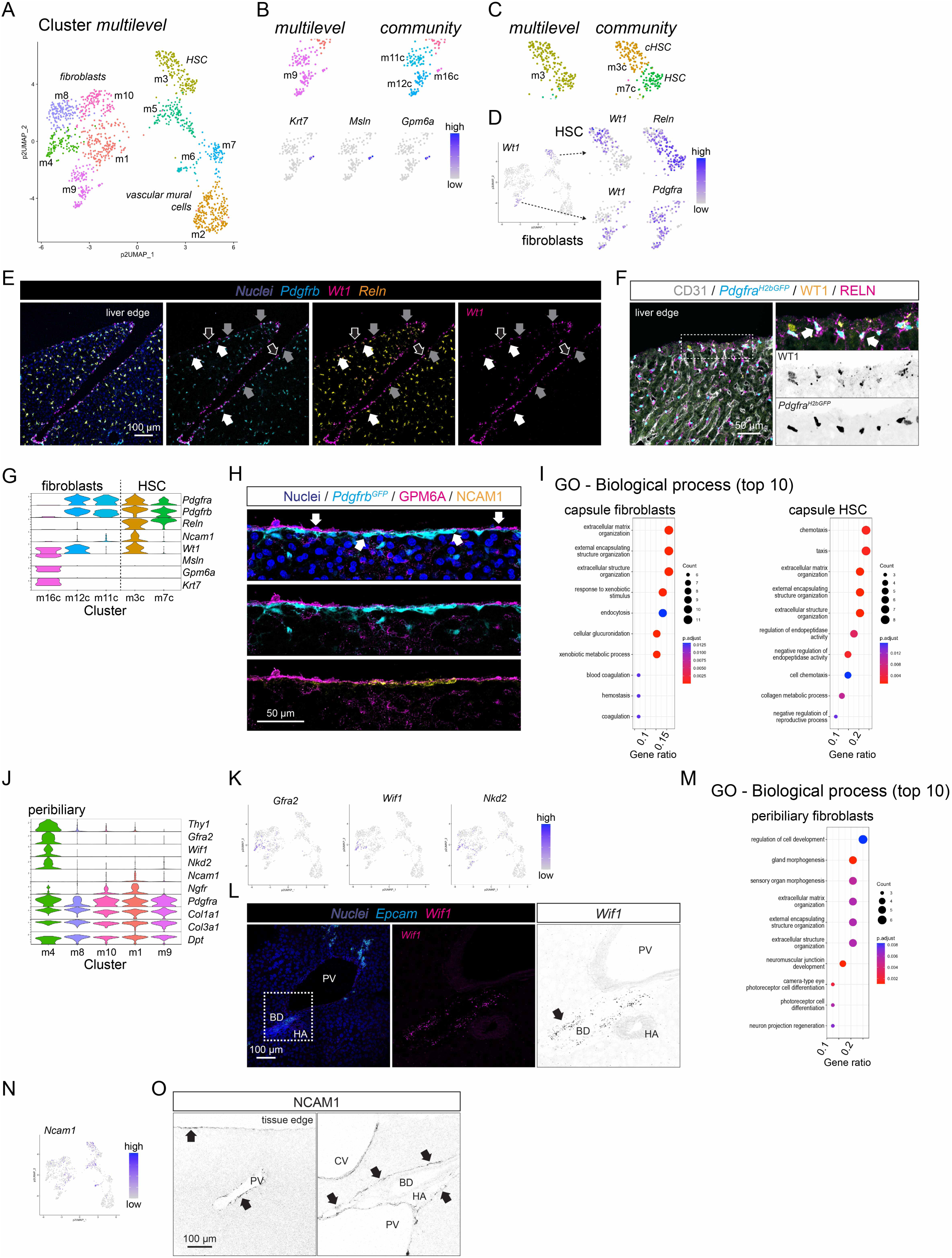
Analysis of mesenchymal cell populations at the liver capsule and portal tract. **A** UMAP visualization of the clustering result for the mesenchymal cell subset. **B** Magnified section of UMAP landscape containing (upper panel) cluster #m9 (multilevel), or clusters #m11c, m12c, and m16c (community), respectively, and (lower panel) UMAP visualization of expression levels of *Krt7*, *Msln*, and *Gpm6a*. **C** Magnified section of UMAP visualization containing cluster #m3 (multilevel), or clusters #m3c, m7c (community), respectively. **D** UMAP visualization of the expression level of *Wt1*, *Reln*, and *Pdgfra* in the mesenchymal cell dataset and magnified sections shown in B and C. **E** ISH for *Pdgfrb*, *Wt1*, and *Reln* on a liver tissue section, focusing on the edge of the tissue (white arrows indicate triple-positive cells, grey arrows indicate *Wt1 Pdgfrb* double positive cells, and open arrows indicate *Wt1* single positive cells). **F** IF for CD31, WT1, and RELN on liver tissue section from a *Pdgfra^H2bGFP^* reporter mouse. The arrows indicate WT1, RELN, GFP triple-positive cells. **G** Violin plot showing the expression of exemplary genes in selected fibroblast and HSC clusters. **H** 3D-redering (angled to achieve 90° view of outer cell layers) of IF with GPM6A and NCAM1 on a liver tissue section from a *Pdgfrb^GFP^* reporter mouse. The arrows indicate the mesothelial cell layer marked by GPM6A and the underlying fibroblast layer indicated by *Pdgfrb^GFP^* and NCAM1. **I** Dot plots showing the top10 overrepresented GO terms of genes with enriched expression in capsular fibroblasts (left panel) or cHSC (right panel). **J** Violin plot showing the expression of genes with enriched expression in the cells from cluster #m4 or cluster #m1 of the mesenchymal cell subset, together with general fibroblast markers. **K** UMAP visualization of the expression levels of *Gfra2*, *Wif1*, and *Nkd2*. **L** ISH for *Epcam* and *Wif1* on a liver tissue section. The arrow highlights *Wif1* expression at the bile duct. **M** Dot plot showing the top10 overrepresented GO terms of genes with enriched expression in peribiliary fibroblasts. **N** UMAP visualization showing the expression level of *Ncam1*. **O** IF for NCAM1 on a liver tissue section, focusing on the edge of the liver (left panel) or the portal tract (right panel). Arrows indicate NCAM1 positive structures. PV: portal vein, BD: bile duct, HA: hepatic artery, CV: central vein. Scale bars are indicated in the respective image panels.

#### Cells at the liver capsule

The liver capsule is known to be covered by an outer monolayer of mesothelial cells (Li *et al*, 2013). In-depth analysis of the mesenchymal cell dataset revealed a small group of mesothelial cells, separated by community clustering (cluster #m16c), and characterized by the expression of *Msln*, *Gpm6a* and *Krt7* (Figure 4B, Figure S4A,B). These cells also expressed the mesothelial cell marker Wilms Tumor 1 transcription factor (encoded by *Wt1*). Notably, *Wt1* was also expressed by subpopulations of fibroblasts and HSC (Figure 4C,D, Figure S4B). Tissue analysis revealed *Wt1*/WT1 expressing cells located at the capsule and sub-capsular region, partially overlapping with expression of *Pdgfrb*, *Pdgfra^H2bGFP^,* and *Reln* (Figure 4E,F, Figure S4C). This suggests the presence of heterogenous fibroblast and HSC subpopulations contributing to the formation of the capsule and sub-capsular region. However, we could also show occasional WT1 *Pdgfra^H2bGFP^* double positive cells in the wall of large central veins (Figure S4C).

Capsular fibroblasts (clusters #m11c, m12c) expressed *Pdgfra* and *Pdgfrb* but not *Reln*, while capsular HSC (cHSC, cluster #m3c) expressed *Reln* and *Ncam1* (encoding the neural cell adhesion molecule 1) in addition to *Wt1* (Figure 4G,H). To gain an overview of the molecular phenotype of capsular fibroblasts and cHSC, we performed DEG analysis comparing the capsular cells to other liver fibroblast or HSC populations, respectively. Capsular fibroblasts displayed distinct expression of genes, such as *Scara5*, *Fbln2*, *Adgrd1*, *Osr1* (encoding the odd-skipped related transcription factor 1), that has been linked to liver fibrosis (Nian *et al*, 2024), as well as mesenchymal collagen production (Murugapoopathy *et al*, 2021) (Figure S4D). In contrast, cHSC expressed many conventional fibroblast markers, including *Gpx3*, *Mfap4*, *Fmod*, and *Dpt*, but also genes not normally associated with fibroblasts, such as *Alcam* (encoding the activated leukocyte cell adhesion molecule) (Figure S4E), which is also expressed by cholangiocytes.

GO analysis of overrepresented genes in capsular fibroblasts or cHSC revealed terms related to ‘extracellular matrix’ and ‘encapsulating’ for both populations, as well as ‘chemotaxis’ specifically for cHSC (Figure 4I). This suggests their active contribution to the formation of the liver capsule and potential roles as progenitors of HSC.

#### Cells at the portal tract and central vein

The portal tract contains the most complex composition of mesenchymal cell types found in the liver. These mesenchymal cells were associated with hepatic arteries, arterioles, portal veins, and bile ducts. We identified fibroblasts residing in the peribiliary niche (cluster #m4) (Figure 4A), characterized by a distinct molecular profile that includes the expression of *Thy1* (encoding CD90), a commonly used fibroblast marker, as well as *Gfra2*, *Nkd2* (encoding the naked cuticle 2), and *Wif1* (encoding the Wnt inhibitory factor 1), the latter two reported as Wnt-pathway antagonists (Gammons *et al*, 2020; Poggi *et al*, 2018) (Figure 4J,K). The localized expression was confirmed by ISH for *Wif1* or *Gfra2* (Figure 4L, Figure S4F). GO analysis of genes overrepresented in peribiliary fibroblasts (Figure S4G) revealed terms related to ‘morphogenesis’ and ‘extracellular matrix’ (Figure 4M). Of note, IF for NCAM1 marked cHSC, but it also highlighted a subset of portal fibroblasts (cluster #m1) that formed a capsule-like structure lining the portal tract (Figure 4N,O). This suggests the presence of heterogenous fibroblast subpopulations contributing to the formation of the portal tract connective tissue and capsule (Glisson’s capsule).

Mural cells distributed to adjacent cell clouds in the UMAP analysis (Figure 5A,B, Figure S5A) and were identified by canonical marker gene expression (Figure 5C, Figure S5B). Notably, scRNA-seq analysis identified *Ncam1* expressing cells amongst the fibroblasts likely involved in the formation of the portal capsule (Glisson’s capsule, see above), and in mural cell clusters (#m5c and m6/m15c), additionally exhibiting expression of *Cdh3*, encoding the cell-cell adhesion protein cadherin 3 (also known as placental (P)-cadherin), and *Npnt* (encoding nephronectin) (Figure 5D,E). Large-caliber portal veins that are muscularized and display a strong and continuous *Acta2*/aSMA signal showed robust IF signals for NPNT (Figure 5F). Scattered NPNT-positive cells were also observed around large and medium-sized central veins (Figure S5C), suggesting that cells in cluster #m6/m15c represent a distinct subpopulation of vascular SMC preferentially found in large-diameter veins.

**Figure 5:**
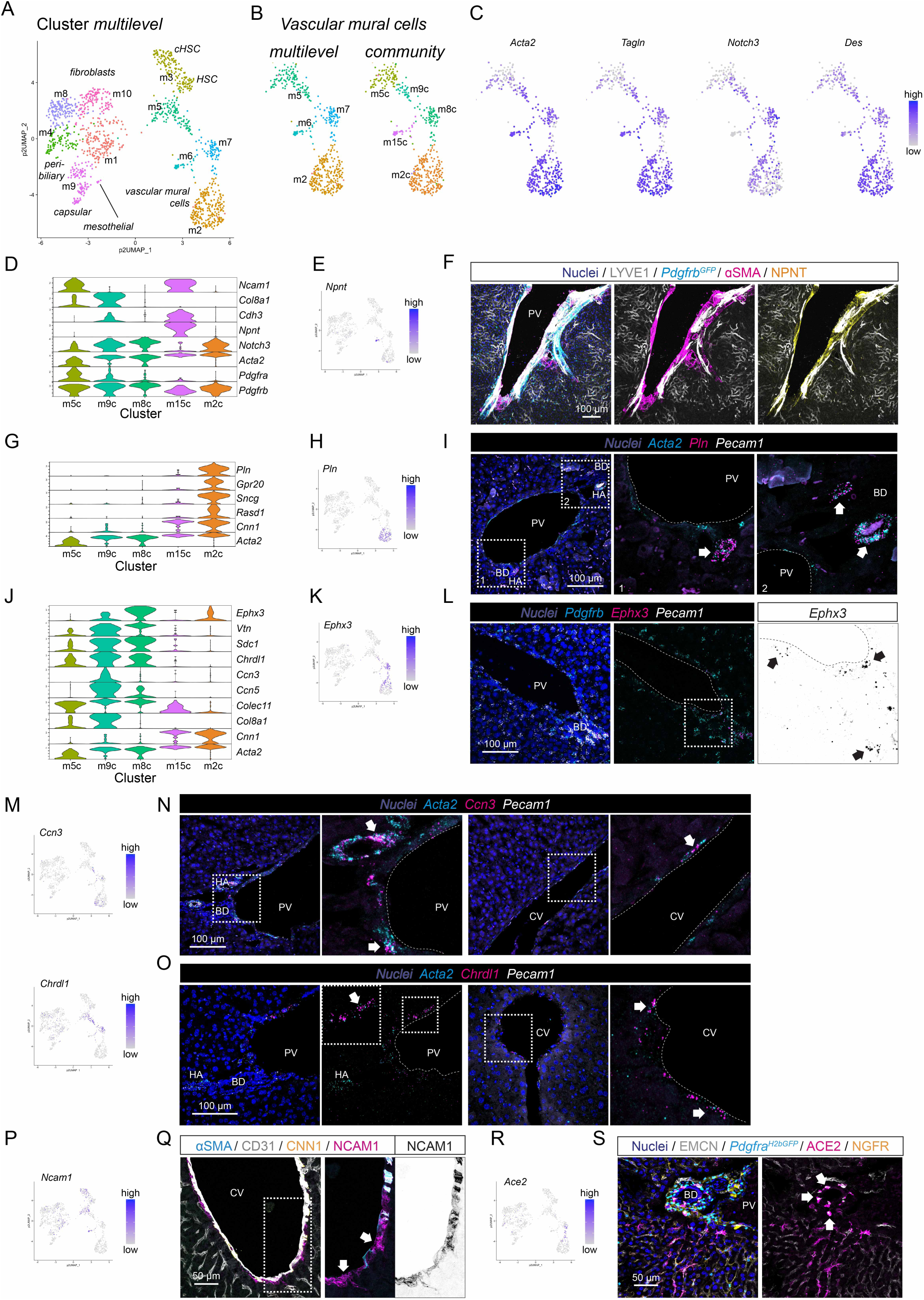
Analysis of perivascular mesenchymal cells in the portal tract and around the central vein. **A** UMAP visualization of the clustering result for the mesenchymal cell subset. **B** Magnified section of UMAP landscape containing vascular mural cell clusters (multilevel #m2, m5, m6, m7; community: #m2c, m5c, m8c, m9c, m15c). **C** UMAP visualization of the expression levels of vascular SMC marker genes (*Acta2*, *Tagln*, *Notch3*, and *Des*) in the magnified section of the UMAP landscape containing vascular mural cell clusters. **D** Violin plot showing the expression levels of exemplary genes differentially expressed between vascular mural cell subpopulations (community clusters). **E** UMAP visualization of the expression level of *Npnt*. **F** IF for LYVE1, αSMA, and NPNT on a liver tissue section from a *Pdgfrb^GFP^* reporter mouse. **G** Violin plot showing the expression level of genes with enriched expression in arterial SMC (cluster #m2/m2c). **H** UMAP visualization of the expression level of *Pln*. **I** ISH for *Acta2*, *Pln*, and *Pecam1* on a liver tissue section. The arrows indicate arterial SMC. **J** Violin plot showing the expression level of genes with enriched expression in venous SMC (clusters #m8c, m9c). **K** UMAP visualization of the expression level of *Ephx3*. **L** ISH for *Pdgfrb*, *Ephx3*, and *Pecam1* on a liver tissue section focusing on the portal tract. The arrows highlight expression of *Ephx3* in the portal tract. **M** UMAP visualization of the expression level of *Ccn3* (upper panel) or *Chrdl1* (lower panel). **N** ISH for *Acta2*, *Ccn3*, and *Pecam1* on a liver tissue section. The arrows indicate *Ccn3* positive SMC. **O** ISH for *Acta2*, *Chrdl1*, and *Pecam1* on a liver tissue section. The arrows highlight *Chrdl1* positive venous SMC. **P** UMAP visualization of the expression level of *Ncam1*. **Q** IF for αSMA, CD31, and NCAM1 on a liver tissue section. The arrows highlight NCAM1 positive αSMA negative cells. **R** UMAP visualization of the expression level of *Ace2*. **S** IF for EMCN, ACE2, and NGFR on a liver tissue section from a *Pdgfra^H2bGFP^* reporter mouse. The arrows indicate ACE2 positive pericytes. PV: portal vein, BD: bile duct, HA: hepatic artery, CV: central vein. Scale bars are indicated in the respective image panels.

The largest group of vascular SMC (cluster #m2/m2c) was characterized by the expression of *Pln* (encoding phospholamban), *Gpr20*, *Sncg*, *Rasd1* and high levels of *Cnn1* (encoding calponin 1) (Figure 5G,H), genes previously reported as arterial SMC markers (Muhl *et al*., 2022b). ISH analysis for *Pln* demonstrated restricted expression in cells of the hepatic artery and larger arterioles around the bile duct (Figure 5I), confirming that the cells in cluster #m2/m2c are SMC from the hepatic arteries and arterioles of the PBV.

Cluster #m7/m8c located adjacent to the arterial SMC and large-diameter vein SMC in the UMAP display and was characterized by the expression of *Ephx3* (encoding the epoxide hydrolase 3) as well as low levels of *Cnn1* (Figure 5J,K). ISH for *Ephx3* revealed specific expression in mural cells of the portal vein and the peribiliary vascular plexus, but not around the central vein (Figure 5L, Figure S5D). We therefore designated the cells in cluster #m7/m8c as portal vein SMC. Cells allocated to cluster #m9c expressed genes previously described as venous SMC markers (Muhl *et al*., 2022b), including *Ccn3* (encoding the cellular communication network factor 3) and *Chrdl1* (encoding chordin-like 1), a BMP-signaling regulator (Figure 5J,M). ISH confirmed the expression of both transcripts at portal and central veins although at lower levels in the central veins (Figure 5N,O). This suggests that cells in cluster #m9c represent a subpopulation of venous SMC distributed throughout the hepatic venous system partially sharing the anatomical niche with vascular SMC of cluster #m7/m8c.

So-called second layer cells (SLC) are morphologically and positionally distinct from HSC and known to reside in the walls of central veins, occupying the space between the central vein SMC and the nearest sinusoids. The number of SLC declines in the walls of smaller central veins, where HSC have been suggested to adopt their position (Bhunchet & Wake, 1992). Cells in cluster #m5c, placed between venous SMC and HSC in the UMAP landscape, expressed *Ncam1* (Figure 5D,P), and IF for NCAM1 showed staining around central veins (Figure 5Q, Figure S5E), and as described above in large muscularized portal veins, the liver capsule, and the portal tract. The NCAM1 staining pattern at central veins, however, aligns with the expected position of SLC, particularly where there is no overlap with αSMA staining (Figure 5Q, Figure S5F). We therefore assume that cluster #m5c represents SLC; yet, the same staining pattern is also observed around portal veins (Figure 5Q, Figure S5E,F), perhaps indicating that SLC may occupy both central and portal areas in the mouse liver. By applying a gene signature that identifies fibroblast-like and mural cell-like expression characteristics (Muhl *et al*., 2020), we found that the SLC cluster #m5c showed greater similarity to fibroblasts than the other mural cell clusters and HSC (Figure S5G).

GO analysis of the different hepatic mural cell populations reveled terms related to ‘muscle tissue development’, ‘serotonin transport’ and ‘neurotransmitter uptake’ associated with large-caliber vein SMC, and terms related to ‘muscle cell development’ and ‘Notch signaling’ overrepresented in arterial SMC. Additionally, terms related to ‘angiogenesis’, ‘artery morphogenesis’ and ‘hepatic stellate cell activation’ were identified for portal vein SMC, while terms associated with ‘extracellular matrix organization’, ‘angiogenesis’ and ‘wound healing’ were noted in venous SMC. For SLC, terms related to ‘extracellular structure organization’, ‘positive regulation of locomotion’ and ‘hormone metabolic process’ were associated (Figure S5H).

Pericytes did not form their own distinct cluster in our dataset, presumably because of their low number. However, cells situated between arterial SMC (cluster #m2/m2c) and portal vein SMC (cluster #m7/m8c) in the UMAP landscape likely represent pericytes of the peribiliary vascular plexus because their expression of *Ephx3*, *Ace2*, *Cspg4* and *Higd1b*, while being negative for *Pdgfra* (Figure 5K,R,S Figure S5J), matches pericytes found in other tissues (Armulik *et al*, 2011; Muhl *et al*., 2020; Vanlandewijck *et al*, 2018).

#### Cells at the sinusoids

HSC reside in the space of Disse between sinusoidal endothelial cells and hepatocytes. HSC have previously been characterized using scRNA-seq, and specific marker genes have been proposed for their identification (Dobie *et al*., 2019; Filliol *et al*, 2022; Krenkel *et al*., 2019; Mederacke *et al*, 2022; Yang *et al*, 2021). With this information as guide, we initially used the expression of *Reln* and *Lrat* to tentatively define HSC clusters (Figure 1L). To achieve a more comprehensive basis for HSC identification, we analyzed earlier Smart-seq2 data from Dobie *et al*. (GSE137720) (Dobie *et al*., 2019) and created an HSC signature comprising 180 genes with enriched expression in HSC (Table 1). The application of the HSC signature score to the mesenchymal dataset revealed cluster #m7c as having the highest score, thus putatively representing HSC (Figure 6A-C, S6A-C). *Ngfr* and *Adamstl2* have been suggested as markers for periportal and pericentral HSC, respectively (Figure S6C) (Dobie *et al*., 2019); however, our transcriptomic data did not identify these subpopulations, likely due to the low number of HSC in the dataset (Figure S6D). Nevertheless, IF analysis confirmed the expected distribution of NGFR-positive HSC towards the portal tract (Figure S6D).

**Figure 6:**
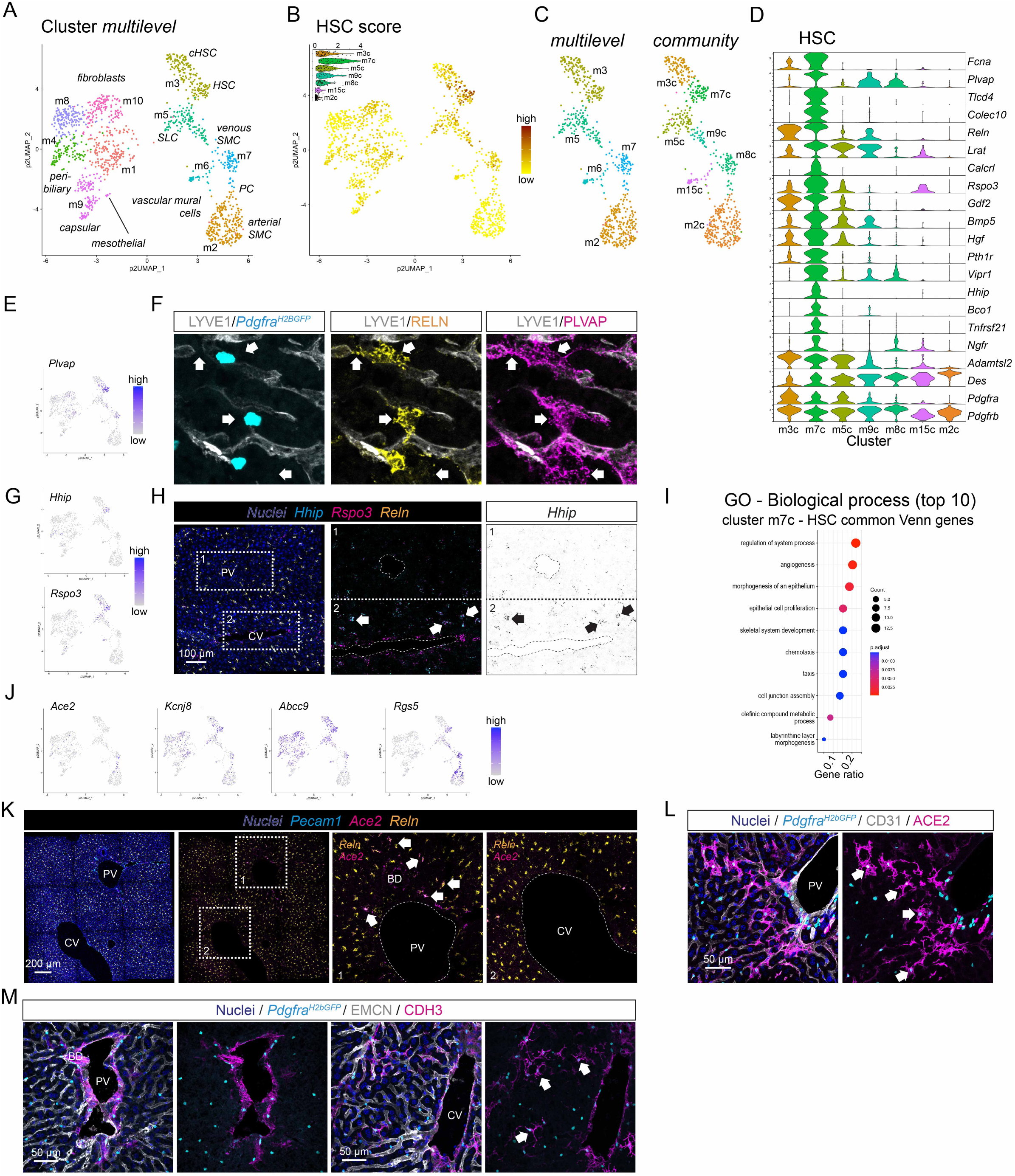
Analysis of mesenchymal cells located at the sinusoids. **A** UMAP visualization of the clustering result for the mesenchymal cell subset. **B** UMAP visualization of the mesenchymal cell subset showing the cumulative expression level of HSC-specific genes (HSC score). The inlayed violin plot shows the individual scores (arbitrary units). **C** UMAP visualization of the magnified section of the UMAP landscape containing HSC and vascular mural cell clusters (multilevel: #m2, m3, m5, m6, m7; community: #m2c, m3c, m5c, m7c, m8c, m9c, m15c). **D** Violin plot showing the expression levels of HSC enriched genes in the HSC and vascular mural cell clusters. **E** UMAP visualization of the expression level of *Plvap*. **F** IF for LYVE1, RELN, and PLVAP in a liver tissue section from a *Pdgfra^H2bGFP^* reporter mouse. Arrows indicate PLVAP RELN double positive HSC. **G** UMAP visualization of the expression level of *Hhip* (upper panel) or *Rspo3* (lower panel). **H** ISH for *Hhip*, *Rspo3*, and *Reln* on a liver tissue section. Arrows indicate *Hhip Rspo3 Reln* triple positive HSC. **I** Dot plot showing the top10 GO terms overrepresented in genes with enriched expression in HSC (cluster #m7c). **J** UMAP visualization of the expression level of pericyte marker genes (*Ace2*, *Kcnj8*, *Abcc9*, and *Rgs5*). **K** ISH for *Pecam1*, *Ace2*, and *Reln* on a liver tissue section. Arrows indicate *Ace2 Reln* double positive HSC. **L** IF for CD31 and ACE2 on a liver tissue section from a *Pdgfra^H2bGFP^* reporter mouse. Arrows indicate GFP ACE2 double positive HSC. **M** IF for EMCN and CDH3 in a liver tissue section from a *Pdgfra^H2bGFP^* reporter mouse focusing on the portal tract (left panel) or central vein (right panel). Arrows indicate GFP CDH3 double positive HSC. PV: portal vein, BD: bile duct, HA: hepatic artery, CV: central vein. Scale bars are indicated in the respective image panels.

**Table 1A:**
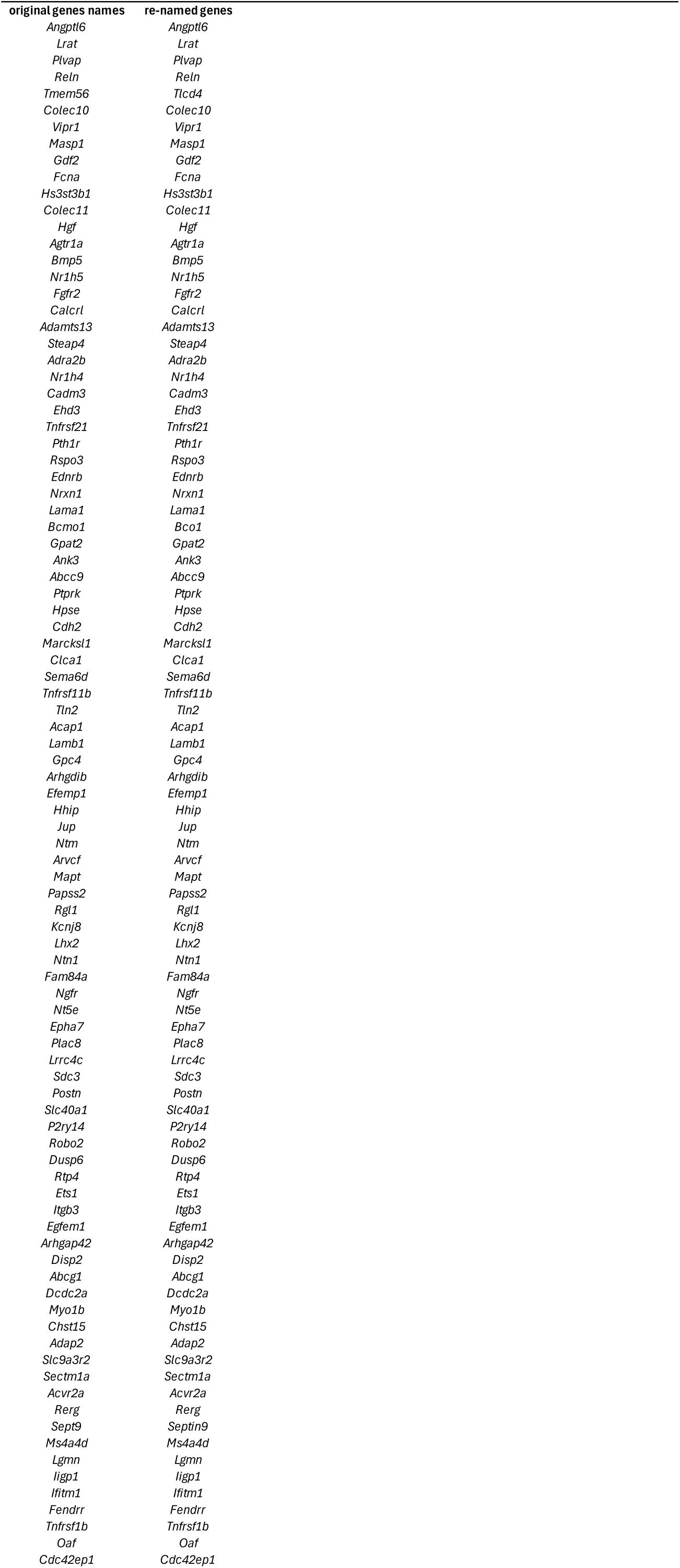
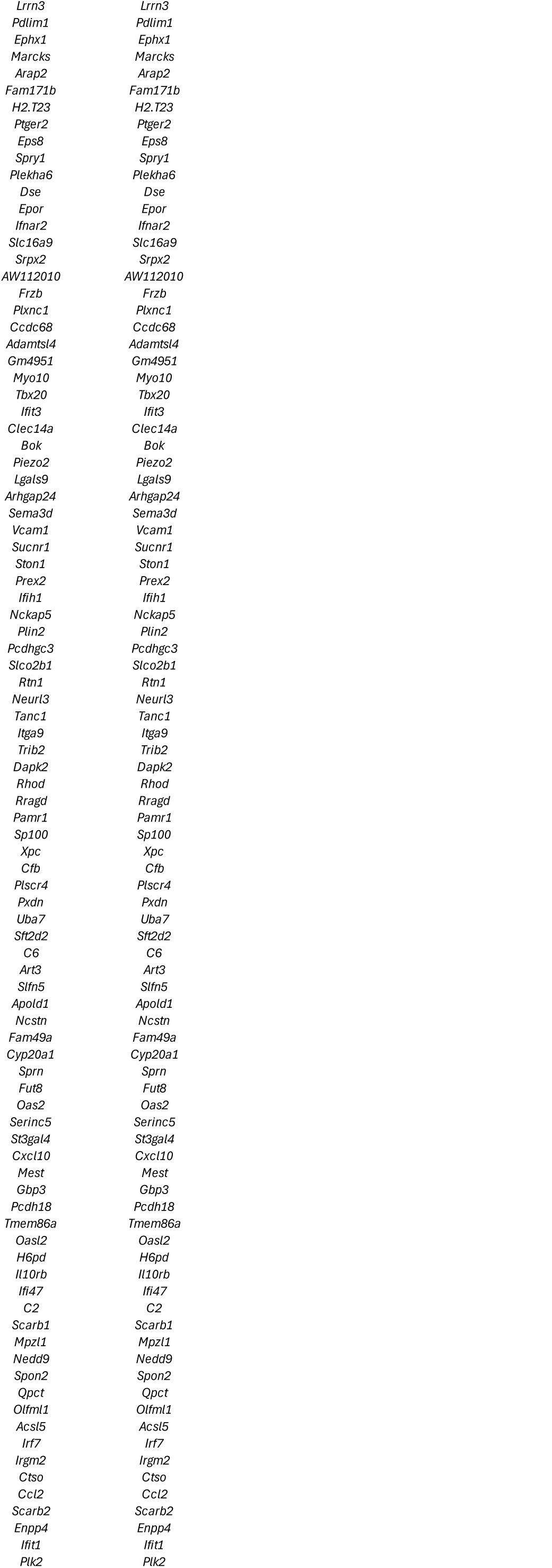
Gene lists of genes with enriched expression in HSC compared with different groups of related cell types.

**Table 1B:**
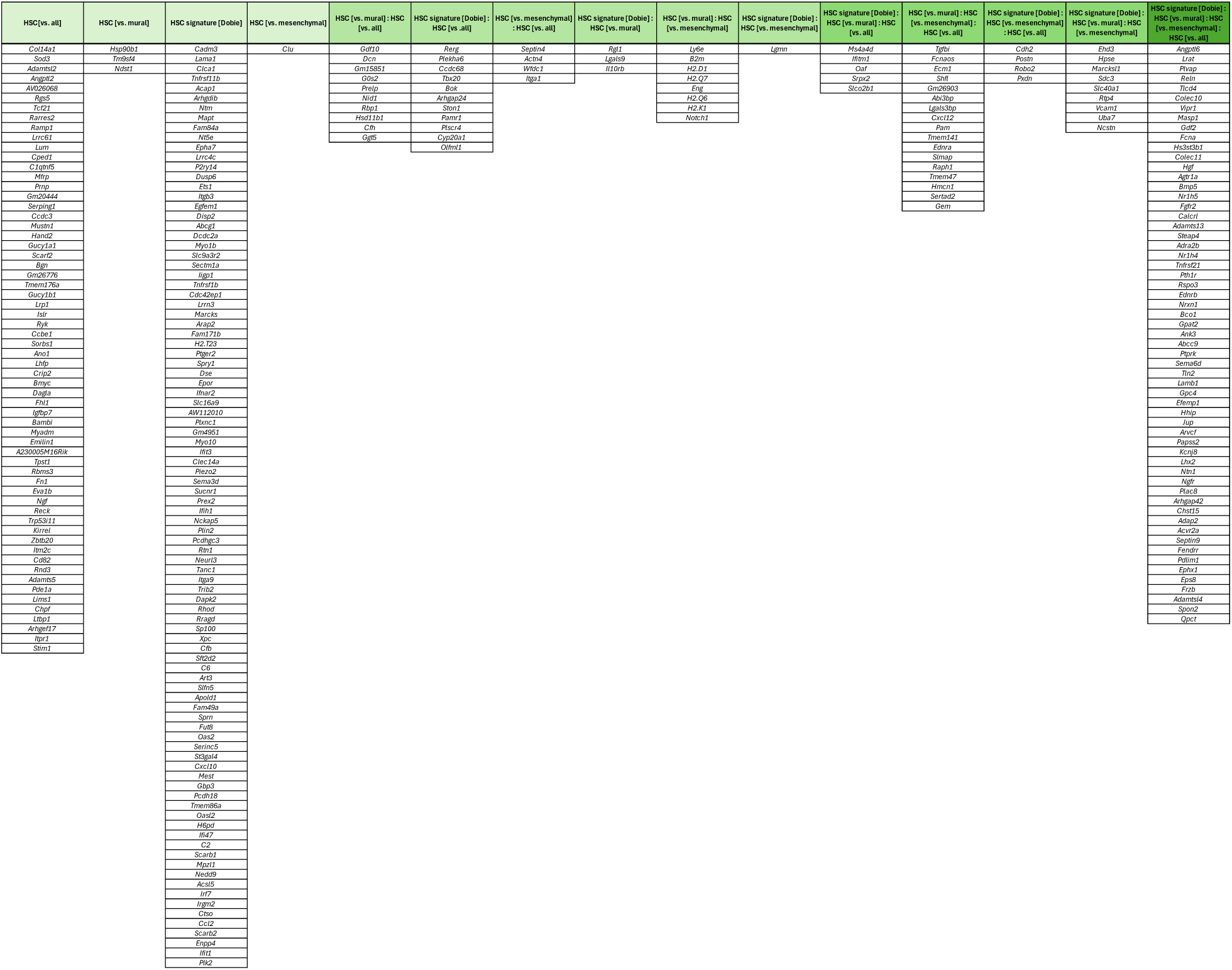
Gene lists of genes with enriched expression in HSC compared with different groups of related cell types.

Through a series of DEG analysis, we generated a core HSC gene set of 59 genes (Figure S6E,F, Table 1). In addition to already established markers, we found that HSC exhibited distinct expression of *Plvap* (encoding the plasmalemma vesicle associated protein, PLVAP), a commonly used endothelial cell marker, *Tlcd4* (previously annotated as *Tmem56*), *Fcna* (encoding ficolin a), *Hhip* (encoding the hedgehog-interacting protein), *Pth1r* (encoding the parathyroid hormone 1 receptor), and *Bco1* (encoding the beta-carotene oxygenase 1) (Figure 6D). IF for PLVAP and RELN confirmed co-expression in *Pdgfra^H2bGFP^* positive HSC (Figure 6E,F, Figure S6G). This finding was further corroborated by ISH for *Plvap* and *Pdgfrb* (Figure S6H,I). HSC localizing to the pericentral region co-expressed *Rspo3* (a marker for central vein EC) and *Hhip* (Figure 6G,H, Figure S6J). GO analysis of the core HSC gene set revealed terms related to ‘angiogenesis’, ‘epithelium morphogenesis’, and ‘olefinic compound metabolism’ associated with HSC (Figure 6I).

Notably, we observed that HSC near the portal tract expressed ACE2 (encoding angiotensin converting enzyme 2), also known as a receptor for SARS-CoV-2 (Muhl *et al*, 2022a), similar to the pericytes of the peribiliary vascular plexus (Figure S6K). The co-expression of ACE2/*Ace2* with HSC markers, such as NGFR (Figure 5S) and *Reln* (Figure 6K), alongside the morphological resemblance to HSC, expression of *Pdgfra^H2bGFP^* (Figure 6L), and the absence of αSMA expression (Figure S6L), suggests that ACE2 marks a subpopulation of periportal HSC. Conversely, a HSC subpopulation localized in the pericentral niche was identified by CDH3 staining (Figure 6M). Like the periportal HSC subpopulation, the pericentral HSC exhibited characteristic HSC morphology and co-expression of CDH3 with *Pdgfra^H2bGFP^* (Figure 6M). However, the portal zonation marker NGFR rarely colocalized with CDH3 positive HSC (Figure S6M,N). Although we could easily identify these two HSC subpopulations through IF analysis, it proved challenging to establish a connection to the transcriptomes in our scRNA-seq dataset. Thus, we concluded that these HSC subpopulations were likely not captured by our scRNA-seq experiments. Examination of the dataset published by Dobie *et al*. demonstrated co-expression of *Ace2* with *Ngfr* and *Myh11* in the HSC cluster, which supported our conclusion. However, *Cdh3* was not expressed amongst the HSC cluster (Figure S6O). Recently, a periportal HSC subpopulation positive for *Myh11* (encoding smooth muscle myosin heavy chain, SMMHC) was identified and suggested to participate in capillarization during hepatic fibrosis (Kan *et al*, 2021), which may reflect the ACE2 positive subpopulation discussed above. Further work is needed to identify the molecular phenotype of the periportal and pericentral HSC subpopulations.

In summary, we present a holistic overview of the complex landscape of mesenchymal cells that populate the various anatomical niches of the adult murine liver. We provide molecular characteristics for capsular, peribiliary, and portal fibroblasts, as well as for arterial and different venous SMC populations. Additionally, we highlight the presence of SLC (which may also be present in the portal vein), canonical HSC, capsular HSC (cHSC), and two rare subpopulations of HSC residing near the portal vein or central vein, respectively.

### Relation to published work and relevance for hepatic diseases

Not all cell subpopulations identified through histological examination were readily identified in our scRNA-seq dataset. To confirm and validate our observations, we therefore expanded our search for the molecular characteristics of the subpopulations missing in our scRNA-seq data by analyzing previously published scRNA-seq datasets from mouse and human liver cells.

#### Hybrid endothelial cells of the portal vein

Portal vein endothelial cells exhibited a hybrid phenotype with features of both venous and arterial endothelium (Figure 2). Although our scRNA-seq dataset did not include true arterial endothelial cells for comparison, we confirmed the hybrid gene expression in portal vein endothelial cells by performing direct comparisons with arterial endothelial cells marked by *SEMA3G* expression in human liver samples (Figure 7A-D, Figure S7A,B) (Buonomo *et al*, 2022; Ramachandran *et al*, 2019). Further, analysis of human liver samples also confirmed the presence of the *CD200*, *RGCC*, *CA4* (human homologue to mouse *Car4*) positive endothelial cell subpopulation putatively originating from the PBV (Figure 7B,D).

**Figure 7:**
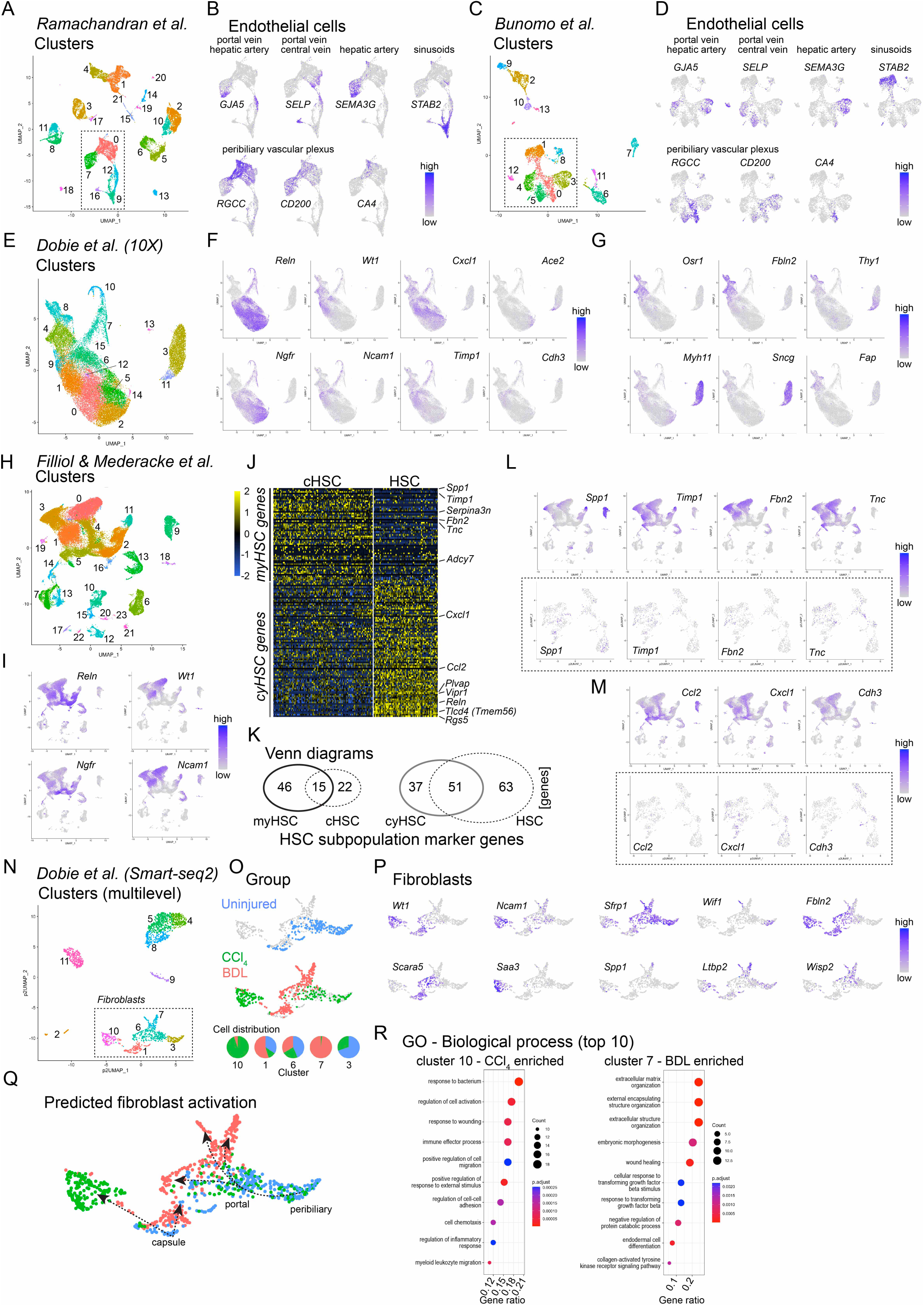
Analysis of previously published scRNA-seq datasets from human and mouse liver tissues. **A** UMAP visualization of the clustering result of the 10X dataset (GSE136103) from Ramachandran *et al*. using human liver cells and Seurat clustering. Clusters containing endothelial cells are indicated (#0, 7, 9, 12, 16). **B** UMAP visualization of the expression level of endothelial cell subtype marker genes (portal vein and hepatic artery, *GJA5*; portal and central vein, *SELP*; hepatic artery, *SEMA3G*; sinusoids, *STAB2*; peribiliary vasculature: *RGCC*, *CD200*, *CA4*) in the magnified section of the UMAP landscape containing endothelial cells clusters. **C** UMAP visualization of the clustering result of the 10X dataset (GSE168933) from Bunomo *et al*. using human liver cells and Seurat clustering. Clusters containing endothelial cells are indicated. **D** UMAP visualization of the expression level of endothelial cell subtype marker genes (portal vein and hepatic artery, *GJA5*; portal and central vein, *SELP*; hepatic artery, *SEMA3G*; sinusoids, *STAB2*; peribiliary vasculature: *RGCC*, *CD200*, *CA4*) in the magnified section of the UMAP landscape containing endothelial cell clusters. **E** UMAP visualization of the clustering result of the 10X dataset (GSE137720) from Dobie *et al*. using all (healthy, CCl_4_ treated for 72h or 6 weeks) cells and Seurat clustering. **F** UMAP visualization of the expression of exemplary mesenchymal cell subset marker genes (*Reln*, *Wt1*, *Ngfr*, *Ncam1*, *Ace2*, and *Cdh3*), or genes with increased expression after CCl_4_ treatment (*Cxcl1* and *Timp1*) in the Dobie *et al*. 10X dataset. **G** UMAP visualization of the expression levels of fibroblast (*Osr1*, *Fbln2*, *Thy1*, and *Fap*) or vascular SMC (*Myh11* and *Sncg*) marker genes in the Dobie *et al*. 10X dataset. **H** UMAP visualization of the clustering result of the combined and integrated 10X datasets (GSE172492 and GSE212047) from Mederacke *et al*. and Filliol *et al*. using all cells (untreated, CCl_4_, HF CDAA, Mdr2KO, and TAZ FPC NASH) and Seurat clustering. **I** UMAP visualization of HSC subset marker genes (*Reln*, *Ngfr*, *Wt1*, and *Ncam1*) in the combined Mederacke *et al*. and Filliol *et al*. dataset. **J** Heap map showing the expression level of the top genes defined by Filliol *et al*. for myHSC or cyHSC, in cHSC or HSC, respectively. Genes with average log-fold ≥1 were selected from the original Supplementary Tables 2 and 3 (Filliol *et al*., 2022). **K** Venn diagrams showing the overlap of top genes defined for myHSC or cHSC (left panel) or cyHSC and HSC (right panel), see also Table 1. **L** UMAP visualization of the expression levels of exemplary genes with increased expression in myHSC in response to pathological challenges (*Spp1*, *Timp1*, *Fbn2*, and *Tnc*) in the combined Mederacke *et al*. and Filliol *et al*. dataset (upper panel), or in the mesenchymal cell subset (lower panel). **M** UMAP visualization of the expression levels of exemplary genes with increased expression in cyHSC in response to pathological challenges (*Ccl2*, *Cxcl1*, and *Cdh3*) in the combined Mederacke *et al*. and Filliol *et al*. dataset (upper panel), or in the mesenchymal cell subset (lower panel). **N** UMAP visualization of the clustering result of the Smart-seq2 dataset (GSE137720) from Dobie *et al*. using all cells (uninjured, CCl_4_ treated, and BDL treated) and pagoda2 multilevel clustering. Clusters containing fibroblasts are indicated (#1, 3, 6, 7, 10). **O** UMAP visualization showing the magnified section of UMAP landscape containing fibroblast clusters color coded for the treatment groups (upper and middle panels), and the percentage of cell distribution per group for each cluster as pie chart (lower panel). **P** UMAP visualization of the expression level of fibroblast subset marker genes (upper panel), or exemplary genes with enriched expression after pathological challenges (lower panel) in the magnified section of the UMAP landscape containing fibroblast clusters. **Q** UMAP visualization showing the distribution of cells color coded for the different treatment groups in the magnified section of UMAP landscape containing fibroblast clusters with putative differentiation paths and fibroblast subset annotated. **R** Dot plot showing the top10 GO terms overrepresented in genes with enriched expression in fibroblasts from CCl_4_ treated (left panel) or BDL treated (right panel) mice, respectively.

#### Rare HSC subpopulations

Periportally and pericentrally localized HSC were clearly identifiable through staining for ACE2 or CDH3, respectively. However, their molecular phenotypes could not be more deeply characterized through our present scRNA-seq data, likely due to their relatively low abundance in the tissue. Despite this, we identified single-cell transcriptomes in 10X Genomics (10X) datasets that were focused to capture HSC from mouse liver samples (Dobie *et al*., 2019) that corroborate the presence of both of these HSC subpopulations (Figure 7E,F, Figure S7C).

#### Mesenchymal cells in hepatic diseases

It is increasingly realized that mesenchymal cells of the liver are important in hepatic disease (Kamm & McCommis, 2022; Wells, 2014a). To gain further insight into the molecular transitions from homeostasis to disease, we incorporated the results from our scRNA-seq analysis with datasets from other sources. To this end, scRNA-seq data from several studies of mouse disease models (Dobie *et al*., 2019; Filliol *et al*., 2022; Mederacke *et al*., 2022) were reprocessed and analyzed. The integrated 10X dataset from Dobie *et al*. (Dobie *et al*., 2019) revealed populations of HSC, actively proliferating HSC, SMC, and fibroblasts (Figure 7E,F, Figure S7C). Notably, a large proportion of HSC originating from injured livers exhibited a transcript pattern similar to cHSC (see below), including the expression of *Wt1*, *Ncam1* and *Cdh3* (Figure 7F, Figure S7C, compare to Figure 4). The fibroblasts displayed a general myofibroblast phenotype (Figure 7G).

Recent scRNA-seq analysis (Filliol *et al*., 2022; Mederacke *et al*., 2022) categorized two distinct HSC subtypes under physiological and pathological conditions: myHSC (myofibroblastic HSC) and cyHSC (cytokine- and growth factor-expressing HSC) (Filliol *et al*., 2022). The integrated analysis of these two studies supported the observation that an HSC population expressing cHSC marker genes (Figure S4E) appears to increase under various pathological challenges (Figure 7H,I, Figure S7D). Comparing the marker genes of myHSC and cyHSC with those of cHSC and general HSC revealed substantial overlap between these subpopulations (Figure 7J,K, Table 2), suggesting that myHSC may originate from cHSC. Considering the gene expression patterns of HSC and cHSC in the homeostatic state, a more refined gene set representing the activation of different HSC subpopulations may be identified (Figure 7L,M). Although we could investigate different HSC populations, the number of fibroblasts in the 10X datasets was limited (Filliol *et al*., 2022; Mederacke *et al*., 2022). Consequently, we analyzed the Smart-seq2 dataset from Dobie *et al*. (Dobie *et al*., 2019), which included transcriptomes from CCl_4_-treated animals and those subjected to bile duct ligation (BDL) (Figure 7N,O, Figure S7E). We identified two specific clusters with strong enrichment of transcriptomes from either CCl_4_-treated samples (cluster #10) or BDL samples (cluster #7) (Figure 7O). Additionally, using the results from our mesenchymal cell dataset analysis, we conducted a preliminary annotation of the fibroblasts of the *Dobie et al.* dataset, which suggested that distinct fibroblast subpopulations, such as peribiliary or capsular fibroblasts, react differently to pathological insults (Figure 7P,Q). GO analysis of enriched genes in the disease-dominant fibroblast clusters revealed terms related to ‘immune process, ‘cell activation, and ‘migration’ to be overrepresented in capsule fibroblasts from CCl_4_-treated animals, whereas terms related to ‘extracellular matrix’, ‘response to transforming growth factor signaling’, and ‘morphogenesis’ were overrepresented in portal and peribiliary fibroblasts from BDL samples (Figure 7R).

In summary, the re-analysis and comparison of previously published scRNA-seq datasets with the data provided herein illustrates the importance of a detailed knowledge of the heterogeneity of HSC and fibroblasts, as identified in our present study, to understand the cell types involved in physiological and pathological processes in the liver.

## Discussion

In the present paper, we provide a comprehensive analysis of hepatic cell types in the adult mouse with particular emphasis on vascular and peri-vascular mesenchymal cells. Our study adds molecular signatures for hepatocytes, cholangiocytes, endothelial cells from the portal vein, sinusoids, central vein, peribiliary vasculature (PBV), and lymphatics, as well as Kupffer cells, and signatures that differentiate between subtypes of hepatic stellate cells (HSC), vascular mural cells, and fibroblasts. Specifically, we outline molecular features to demarcate up to four subtypes of HSC, several different fibroblast populations, and substantial heterogeneity among vascular mural cells, including vascular SMC that form the walls of portal and central veins. Our data can be explored in an appended database, providing online access to gene-by-gene expression patterns at: https://betsholtzlab.org/Publications/LiverScRNAseq/search.html.

Additionally, we provide examples of how our refined liver cell type annotation can be used together with previously published scRNA-seq data from human and mouse samples in order to verify the hybrid nature of portal vein endothelial cells in the human liver and illustrate the importance of well-defined cell type classification at the homeostatic state when inferring phenotypic changes in distinct cell types in response to pathological challenges.

Mesenchymal cells in the liver, in particular HSC, become activated during liver fibrosis. This can lead to the production and deposition of high amounts of extracellular matrix components, which ultimately form a fibrotic scar that impedes liver function (Koyama & Brenner, 2017; Mederacke *et al*, 2013). But the origin of these reactive HSC during liver fibrosis remains enigmatic. We describe a population of *Wt1*-expressing HSC (cHSC) located below the liver capsule, which has been suggested as the source of HSC populating the entire liver (Asahina *et al*, 2011; Ijpenberg *et al*, 2007). Analysis of published scRNA-seq studies indicates that this *Wt1*-positive population expands in models of liver injury and disease. While mesothelial-mesenchymal transition has been suggested to account for the emergence of *Wt1*-positive HSC in disease (Li *et al*., 2013), we demonstrate that a *Wt1*-positive HSC population (cHSC) exists already during homeostasis and may therefore be the source of fibrogenic cells during liver disease. Based on the revised cell type definitions presented herein, lineage tracing studies (past and future) should be reevaluated to clarify the origin of myofibroblasts (assumed to be activated HSC), i.e. whether they differentiate from *Wt1*-positive cHSC, arise from the de novo differentiation of mesothelial cells, or derive from activated HSC from periportal, midlobular, or pericentral location. Additionally, the possible contribution of *Wt1*-positive capsular fibroblasts requires clarification through further studies.

The heterogeneous nature of portal vein endothelial cells––exhibiting arterial and venous features––has, to the best of our knowledge, so far not been described in vascular beds of other organs and likely reflects the unique physiological function of portal veins that conduct blood to capillary beds, in contrast to other veins that direct blood back to the heart. Notably, a similar arterial/venous hybrid phenotype was not noticed in portal vein mural cells, which exhibit molecular phenotypes resembling those in venous mural cells of other tissues (Muhl *et al*., 2022b). We identified several subtypes of venous SMC in the walls of hepatic veins. Portal veins exhibited a higher number of SMC and greater variety of venous SMC subpopulations, compared to central veins, likely reflecting the higher-pressure profile for the portal venous system. The underlying cause and function of portal vein endothelial hybridism requires further investigation to disentangle developmental cues, the physiological, and potential pathological roles of this specialized endothelial cell subpopulation.

The presence of KIT positive sinusoidal endothelial cells in the terminal sinusoids but not in the central vein endothelial cells suggests a distinct function of the sinusoids at this location, which however remains unclear. Notch-signaling has been shown to regulate the expression of *Kit* in sinusoidal endothelium (Duan *et al*., 2022), and in a loss-of-function model for *Dll4*-*Notch1* signaling, it was demonstrated that pericentral sinusoidal endothelial cells were the most reactive endothelial population (Fernandez-Chacon *et al*, 2023), despite a low endogenous expression of *Dll4*.

The PBV arises from the hepatic artery and supports cholangiocyte function (Gaudio *et al*, 1996; Haratake *et al*, 1990). Fine-tuned paracrine signaling between cholangiocytes and vascular cells of the PBV regulates angiogenesis and their morphology. Capillary endothelial cells of the PBV feature distinct molecular phenotypes compared to hepatic sinusoids and are further supported by mural cells that resemble pericytes of other peripheral organs, such as the heart. Capillary endothelial cells of the PBV express genes indicative for fenestrations, suggesting a potential exchange of macromolecules between the blood and biliary cells (Morell *et al*., 2013). The role of vascular and perivascular cells of the bile duct system in portal and biliary fibrotic processes (Wells, 2014b) warrants further dedicated analysis. Such approaches can now be undertaken using cell type specific markers described herein. For example, the lack of cholangiocyte-specific promoters has hampered the progress in the understanding of cholangiocyte physiology and pathology (Banales *et al*., 2019), and our transcriptomic characterization of these cells should aid the development of new genetic tools for the study of cholangiocyte biology. This includes techniques, such as Cre-recombinase-mediated cellular labeling and targeting of cholangiocytes using transgenes driven by hepatic cholangiocyte-specific (among liver cells) promoters, for example *Pkhd1*, *Slc5a1*, or *Cldn8*.

Kupffer cells are the resident macrophage population of the liver, patrolling between sinusoidal lumen and space of Disse (Guilliams & Scott, 2022). A unique feature of Kupffer cells among macrophages is the expression of CDH5 (VE-cadherin) (Scott *et al*, 2018), a commonly considered endothelial cells-specific protein, which has recently been identified also in fibroblast populations of the mouse meninges (Mapunda *et al*, 2023; Pietila *et al*, 2023). The functional consequence of CDH5 expression by Kupffer cells remains unknown, but one can speculate that homotypic interaction of CDH5 aids cell-cell contacts between Kupffer cells and sinusoidal EC. Homodimers of CDH5 link adjacent endothelial cells together, which is important for endothelial barrier integrity, intercellular communication, and contact inhibition (Giannotta *et al*, 2013). Our analysis demonstrated distinct staining patterns for CDH5 between inter-sinusoidal endothelial cell contacts (linear) or sinusoidal endothelial cell-Kupffer cell contacts (patchy). Such ‘patchy’ CDH5 signal has been demonstrated to occur in endothelial cells that actively migrate and was designated as JAIL (junction-associated intermittent lamellipodia) (Cao *et al*, 2017). Another study demonstrated that CDH5 is required for directional migration of endothelial cells (Tonami *et al*, 2023). Whether the expression of CDH5 by Kupffer cells facilitates their migratory capacity along the sinusoidal network remains to be clarified.

In conclusion, this comprehensive atlas of murine liver cell types will be useful––and hopefully inspire new research––to further explore the various roles of distinct hepatic cell types during health and disease.

### Limitations of the study

Although we provide transcriptomic and histological analysis to demarcate more than 20 different cellular subpopulations of the adult mouse liver in homeostasis, we cannot formally exclude the possibility that other resident cell types have been lost during the isolation protocol. For example, two populations of cholangiocytes (large and small) have been described (Tabibian *et al*., 2013), while in our dataset the transcriptomes of cholangiocytes appeared homogenous. Another example is the absence of hepatic artery endothelial cells in our dataset, despite efficient capture of arterial SMC. We have utilized transgenic reporter animals for targeted sorting of cells and cannot exclude the possibility that expression of the fluorescent proteins may have inflicted toxicity and consequently altered the transcriptome of the cells. The observed–– previously unexpected––heterogeneity of mural cell, HSC, and fibroblast populations led to comparably small cell numbers for certain subpopulations (also for endothelial cells of the PBV). However, the Smart-seq2 platform provides deep sequencing reads, that allows for cell type annotation from small numbers of cells. Further, employing IF and ISH analysis we identified two distinct HSC populations that reside close to the portal tract or central vein, respectively. These HSC subpopulations were not captured for our single-cell transcriptome profiling, however, with the herein provided information, dedicated approaches for the characterization of these rare HSC populations can be designed.

All studies of this type have inherent limitations that include the process of cell type annotation and the criteria for inclusion into or exclusion from pre-existing cell classes. The liver mesenchyme represents a good example for this problem–– exhibiting substantial overlapping expression patterns for many commonly applied marker genes (e.g., *Ngfr*, *Reln*, *Lrat*, *Pdgfra*, *Pdgfrb*, *Acta2*, *Kcnj8*, *Abcc9*). Consequentially arising questions are; Where should one separate and demarcate HSC from SLC or fibroblasts? When does one annotate a mural cell as SMC, a SMC subtype, a pericyte, or a SLC? The simple use of the cluster assignment as an unbiased classification is tempting, however it inherits the problems that the results are dependent on the set parameters of the algorithms (e.g. the *multilevel* or *community* setting in pagoda2) and the overall heterogeneity of the input data. Presumably, there is no golden rule that fits all single-cell datasets and types of analysis. Researchers are required to test for and apply the best compromise for each dataset in question to obtain the most meaningful output.

For validation and to link the annotation of transcriptomes to anatomic locations we used image analysis of IF, ISH, and transgenic reporters. Although this is essential, these procedures are biased and dependent on the quality and availability of the reagents. For example, to pinpoint the anatomic location of capsular fibroblasts we used the most relevant markers obtained from the scRNA-seq analysis (*Scara5*, *Osr1*), however, we did not find a clear staining pattern for these markers at the liver capsule, or elsewhere in the tissue. Therefore, we relied on fewer markers to annotate capsular fibroblasts (*Wt1 Pdgfra* double-positive, *Reln*-negative), which however does not exclude the possibility that cells here annotated as capsular fibroblasts may also locate to other anatomical locations.

Our choice of ISH probes and IF antibodies to detect different genes/proteins to identify specific cell subpopulations was done to the best of our judgment, although, new and improved reagents may allow for more refined analysis in the future. Similarly, new iterations of genetic lineage-tracing and reporter models, possibly created on the information from this and/or other contemporary studies may generate unambiguous results and interpretations that allow for further revised cell type identification.

## Methods

### Animals

All experimental procedures involving animals were carried out in accordance with the Swedish legislation and local regulations and guidelines for animal welfare. All mouse experiments were approved by local authorities: Linköpings Animal Research committee – ID 729, 3711-2020, and 11939-2023, and Uppsala Animal Research committee – approval ID 03029/20290. Mice were housed in standard, single ventilated cages with 12h light, 12h dark cycle, ad libitum access to water and chow and an ambient temperature of 20 ± 2°C with a relative humidity of 50 ± 5%. The following mouse strains were used in the study: *C57BL6/J* (The Jackson Laboratory, *C57BL6/J*, were maintained as breeding colony at the local animal facility), *Pdgfrb^GFP^* (Gensat.org, Tg(*Pdgfrb*-eGFP)JN169Gensat/Mmucd), *Pdgfra^H2bGFP^*(*Pdgfra^tm11(EGFP)Sor^* (Hamilton *et al*, 2003), a gift from P. Soriano), *Cspg4^dsRED^* (The Jackson Laboratory, Tg(*Cspg4*-DsRed.T1)Akik/J), *Acta2^GFP^* (The Jackson Laboratory, Tg(*Acta2*-GFP)1Pfk) (Yokota *et al*, 2006), *Cldn5^GFP^*(Tg(*Cldn5*-GFP)Cbet/U), and combinations of these strains. All reporter transgenes were kept as heterozygous. For experiments, adult male and female mice with an age between 8 weeks and one year were used. Liver tissue samples from *Gja5^GFP^* mice (*Gja5^tm1Lumi^*) (Miquerol *et al*., 2004), were generously provided by L. Miquerol and R. Kelly (IBDM, Aix-Marseille University, France).

### Preparation of single-cell solutions from mouse liver tissue

Mouse liver tissues were handled and processed using the same protocol as described before (Muhl *et al*., 2020; Muhl *et al*., 2022b): In brief, animals were euthanized by cervical dislocation and the liver was dissected out and placed into ambient-tempered phosphate-buffered saline (PBS) solution (DPBS, ThermoFisher Scientific), until further processing. The liver tissue was then cut into smaller pieces with scissors and scalpel and incubated in dissociation buffer (Skeletal muscle dissociation kit from Miltenyi Biotec, supplemented with 1 mg/ml Collagenase IV type-S from Sigma-Aldrich) at 37°C with orbital shaking at 500-800 rpm. Mechanical disintegration by pipetting was applied to facilitate tissue disintegration in three to four cycles with 10 min incubation intervals. After dissociation, the remaining tissue debris were removed by sequential passing of the cell solution through a 70 µm and a 40 µm cell strainer. The 70 µm strainer was additionally washed with 5 ml of DMEM (ThermoFisher Scientific) to recover cells adherent to the surface. Thereafter, the cells were pelleted by centrifugation at 300 x g for 5 min. The supernatant was removed, and the cell pellet resuspended in FACS buffer (DPBS supplemented with 0.5% bovine serum albumin, 2 mM EDTA, 25 mM HEPES). The cell suspension was then labeled with different combinations of fluorophore-conjugated antibodies (anti-CD31, anti-EPCAM, anti-CD68, anti-PDGFRα, anti-VE-Cadherin) to mark distinct cell populations for capture (Table S1) for 20-30 min at room temperature (RT). Thereafter, the cells were pelleted by centrifugation at 300 x g for 5 min, the supernatant removed, and cells resuspended in ice-cold FACS buffer and placed on ice until further processing.

### Fluorescent activated cell sorting (FACS)

Single-cell analysis, selection, gating, and deposition of selected droplets (single cells) was done as described before (Muhl *et al*., 2020; Muhl *et al*., 2022b). The antibody stained single-cell suspensions were analyzed using a FACSAria III or FACSMelody (Becton Dickinson Biosciences) cell sorter, each equipped with a 100 µm nozzle. Single-cell events meeting the criteria as described below were collected by deposition into 384-well plates containing 2.3 µl lysis buffer (0.2% Triton X-100, 2 U/ml RNase inhibitor, 2 mM dNTP, 1 µM Smart-dt30VN, ERCC at 1:4 x 10^4^ dilution) per well. Of note, the analysis of single-cell events at the cell sorter was not the basis for cell type identification, but for the enrichment of target cell populations dependent on the signal of expressed reporter genes or antibody labeling. For sorting into 384-well plates, first debris and red blood cells were excluded by setting a generous gate with forward scatter-area/ side scatter-area (FCS-A/SSC-A, linear scale). For doublet discrimination, a second gate using SSC-A/SSC-height and FSC-A/FSC-height was used. Thereafter cells were analyzed for their fluorescent signals and separated dependent on their reporter gene expression and antibody labeling. Fluorescent signals were controlled applying the “fluorescence minus one” method, using samples without antibody labeling and/or from reporter gene negative animals.

Endothelial cells were collected on their labeling with antibodies against CD31 and/or VE-Cadherin. Of note, we noticed that sinusoidal endothelial cells exhibited a lower staining intensity for CD31 but higher staining for VE-Cadherin, compared to endothelial cells from the portal and central veins (CD31^high^ VE-Cadherin^low^), or from *Cldn5^GFP^* mice dependent on the reporter gene signal. Epithelial cells (cholangiocytes) were selected dependent on their anti-EPCAM antibody labeling. Macrophages, targeting resident Kupffer cells, were collected dependent in their anti-CD68 antibody labeling. Hepatic stellate cells, fibroblasts, pericytes, smooth muscle cells and other vascular mural cells were sorted dependent in their distinct *Pdgfra^H2bGFP^*, *Pdgfrb^GFP^*, *Acta2^GFP^* reporter gene expression patterns as well as anti-PDGFRα antibodies. Hepatocytes were collected by unbiased sorting a small amount of living cells. Before sorting, the 384-well plates containing lysis buffer were briefly centrifuged to ensure the proper dispersion of the lysis buffer in the bottom of the wells. Notably, the correct deposition of the selected droplets (i.e., single cells) was controlled by test-spotting (aiming) of beads or cell populations from control samples onto the seal of the respective 384-well plate. If needed, the plate holder position was adjusted for a centered deposition of the droplets. This procedure was performed for each individual plate. The sample-stand and plate-holder were maintained at 4°C during the analysis and sorting procedure. Finished sorted plates were immediately sealed and placed on dry-ice and stored at -80°C, until further processing.

### scRNA-seq library preparation and sequencing

Library preparation and sequencing was performed as described before (Muhl *et al*., 2020; Muhl *et al*., 2022b), and according to the previously established protocol for Smart-Seq2 (Picelli *et al*., 2014). In brief, poly-adenylated mRNA was transcribed to cDNA using oligo(dT) primer and SuperScript II reverse transcriptase (ThermoFisher Scientific). Second strand cDNA synthesis was achieved using a template switching oligo, and the double stranded cDNA was then amplified with PCR. Amplified cDNA was purified, and quality was controlled (QC) by analyzing randomly selected wells (single cell samples) on a TapeStation 4200 or a 2100 Bioanalyzer using DNA high sensitivity chips (Agilent Biotechnologies). When samples (plates) passed the QC, the cDNA was enzymatically fragmented and tagged using Tn5 transposase. Each single well was uniquely indexed using the Illumina Nextera XT index kits (set A-D). Finally, the uniquely indexed cDNA libraries from one 384-well plate were pooled and sequenced together on one lane of a HiSeq3000 sequencer (Illumina), using the sequencing strategy of dual indexing and single 50 base-pair reads.

### Sequence data processing

The obtained sequences (as outlined above) were handled for demultiplexing, mapping and generation of per cell and gene raw-count expression matrices as described earlier (Muhl *et al*., 2020). ENSEMBLE identifiers were annotated using the org.Mm.eg.db package (version 3.14.0) in R-software, retaining ERCC counts in the expression matrix as technical control. Annotated raw counts were loaded into the Seurat package (version 4.3.0) (Satija *et al*., 2015). Low-quality cells (≤ 50 000 counts library size, ≤ 1500 expressed genes, ≥ 10% mitochondrial genes, ≥ 10% ERCC counts), as well as putative doublets (≥ 12 000 expressed genes) were removed from the dataset. Low expressed genes: expressed in ≤ 3 cells with a detection limit = 20 counts per gene, and ≤ 300 total counts per gene, were also removed from the dataset before further processing. Additionally, cells that showed a transcriptome with clear signs of cross cell type contamination were removed from the dataset. After filtering, the dataset consisted of in total 3,491 single-cell transcriptomes, collected from 18 individual mice.

Smart-Seq2 data from this and earlier studies (see also below) were processed using the pagoda2 (version 1.0.10, https://github.com/hms-dbmi/pagoda2) R-software package (Fan *et al*., 2016). General attributes, such as PCA (nPCS = 50, n.odgenes = 3000) and nearest neighbor clustering were performed using default parameters of pagoda2 (k = 30, distance = “cosine”). Dimensional reduction analysis was done using the UMAP function (uniform manifold approximation and projection, n.neighbors = 30, metric = “cosine”, dims = 1:50). Pagoda2 clustering results using the multilevel as well as community setting were considered for downstream analysis. Pagoda2 results were stored within the Seruat object and data visualizations were prepared using functions of the Seurat R-software package (dot plot: DotPlot(), UMAP: DimPlot() or FeaturePlot(), violin plot: VlnPlot(), heatmap: DoHeatmap), as well as the R-software package pheatmap (version 1.0.12) for heatmaps, and clusterProfiler (version 4.2.2) for dot plots of gene ontology (see below) results. For the analysis of selected cell type datasets (parenchymal, endothelial, immune, mesenchymal) the same procedure as described above was applied. For the construction of the bar plot graphs the pagoda2 clustering results were used as a basis. Dependent on the resolution, clusters defined by the multilevel setting, or community setting were used. The order of the clusters within the bar plot graphs was manually defined. The distribution of cells within each cluster was calculated using the SPIN algorithm (/backspin -i input.cef -o output.cef -f 1000 -b both) (Tsafrir *et al*, 2005). For the bar plot graphs, the library size for each cell was normalized to 500 000 counts.

For differential expressed gene (DEG) analysis the FindMarkers() function of the Seurat R-software package was applied, using the MAST test (MAST R-software package, version 1.20.0) for adjusted p-value calculation. Thresholds used for gene-qualification from DEG analysis were an adjusted p-value ≤ 0.05, the expression in ≥ 30% of cells and a log_2_ fold-change ≥ 0.5 or 1 in the respective group.

### Gene ontology (GO) analysis

For gene ontology (GO) analysis the clusterProfiler (version 4.2.2) R-software package was used (Wu *et al*, 2021). In brief, the enrichGO() function was used to identify enriched GO terms from the Biological Process subontology. The list for all genes in the respective dataset was used as reference (gene universe). Terms with an adjusted p-value ≤ 0.05 (pvalueCutoff = 0.05, pAdjustedMehod = “BH” (BH = Benjamini-Hochberg)) were selected and after application of the simplify() function (cutoff = 0.7), to omit redundant terms, the top 10 terms (min-GSSize = 10, maxGSSize = 500) were displayed in the respective dot plots, created with the dotplot() function of the clusterProfiler R-software package.

### Published scRNA-seq datasets available in the public domain

Single-cell transcriptomes from earlier studies of human and mouse liver tissues were re-processed and analyzed in the study. We acquired the deposited raw data from the NCBI gene omnibus database with the following accession numbers: GSE158183, GSE172492 (Mederacke *et al*., 2022), GSE212047 (Filliol *et al*., 2022), GSE168933 (Buonomo *et al*., 2022), GSE136103 (Ramachandran *et al*., 2019), and GSE137720 (Dobie *et al*., 2019).

Processing and analysis of earlier Smart-Seq2 datasets was done as described above. The HSC signature was calculated from the GSE137720 dataset, using only cells from uninjured samples. Differential gene expression was performed as described above and genes highly enriched in HSC clusters, compared to all other cells were used to determine the signature score for our dataset.

Processing and analysis of 10X datasets was done using the Seurat R-software package. In brief, raw count data was loaded into a Seurat object and low-quality cells were removed (≤ 500 detected genes, ≥ 10% counts from mitochondrial genes). The default Seurat pipeline was used to calculate general dataset features, such PCA, clusters, and UMAP distribution. When several samples were combined for analysis, but showed sample-specific clustering, technical batch effects were assumed, and data-integration functions of the Seurat R-software package (SelectIntegrationFeatures(), FindIntegrationAnchors(), IntegratData()) were applied before resuming the analysis of the dataset. Samples from GSE158183, GSE172492 and GSE212047, are thematically related (Filliol *et al*., 2022; Mederacke *et al*., 2022), were integrated, and analyzed together. Data visualization and differential gene expression analysis was performed using functions implemented in the Seurat R-software package as described above.

### Immunofluorescence staining

Standard procedures for immunofluorescence (IF) staining were applied. In brief, liver tissues were harvested from euthanized mice as described above and, if not otherwise stated, immersion fixed using 4% buffered formalin solution (Histolab) for 4 – 12h at 4°C. After fixation, the tissue samples were immersed in 20 – 30% sucrose/PBS solution for at least 24h at 4°C before further processing. For cryo-sections, tissues were, if necessary, carefully trimmed and dissected and then placed in cryo-molds and embedded using NEG50 cryo-medium and sectioned on a Cryostat NX70 (ThermoFisher Scientific) or Cryostat (Leica) into 14 – 30 µm thick sections, collected on SuperFrost Pluss glass slides (Metzler Gläser). Sections were stored at -20 to - 80°C until further processing.

For staining, sections were placed at RT and allowed to dry for 15 min. Thereafter, the sections were briefly washed in PBS and then incubated with blocking buffer (Serum-free protein blocking solution, DAKO), supplemented with 0.2% Triton X-100 (Sigma-Aldrich). After the blocking, the sections were sequentially incubated with primary antibodies, diluted in blocking buffer supplemented with 0.2% Triton X-100, and corresponding fluorophore-conjugated secondary antibodies, diluted in blocking solution, according to the manufacturers’ recommendations (Table S1). For nuclear staining, Hoechst 33342 (trihydrochloride, trihydrate, ThermoFisher Scientific) at 10 µg/ml was added to the secondary antibody solution. Thereafter, sections were mounted using ProLong Gold® mounting medium (ThermoFisher Scientific) and sections stored at 4°C. Micrographs were acquired using a Leica TCS SP8 confocal microscope with LAS X software (version 3.5.7.23225, Leica Microsystems), or a Nikon Eclipse Ti2 confocal microscope, equipped with iXon EMCCD and iZyla SCMOS cameras, with Fusion software (version 2.4, Dragonfly 505 high speed confocal platform, Andor Technologies, Inc). The acquired images were graphically processed and adjusted individually for brightness and contrast using ImageJ/FIJI software (Schindelin *et al*, 2012) for optimal visualization. All images, if not otherwise stated, are presented as maximum intensity projections of the acquired z-stacks, covering the entire thickness of the respective sections.

### In situ hybridization staining (RNAscope®)

For fluorescent, multiplexed in situ RNA hybridization (ISH) the RNAscope® Fluorescent Multiplex Assay system (Advanced Cell Technologies), without or with TSA-amplification (V2 kit) was applied according to the manufacturers’ recommendations. Cryo-sections from liver tissue were prepared as described above (Immunofluorescent staining), with or without (fresh frozen) immersion fixation before sectioning. After dehydration, the sections were prepared using Pretreat 4 solution for 15 – 30 min at RT. RNAscope® probes (Table S1) were applied according to the manufacturers’ recommendations and incubated at 40°C for 2h. After completion of the protocol as described by the manufacturer, the sections were mounted using ProLong Gold® mounting medium and stored at 4°C. Micrographs were acquired as described above (Immunofluorescent staining).

### Statistics and reproducibility

Gene ontology (GO) analysis was performed using the clusterProfiler R-software package (Wu *et al*., 2021). Term enrichment was considered significantly when the adjusted p-value, using the Benjamini-Hochberg procedure, was ≤ 0.05. All immunofluorescence (IF) and RNA in situ hybridization (ISH) experiments were performed at least twice using the same or varying antibody, or probe combinations, respectively. Tissue sections from at least two individual mice were analyzed for each respective IF or ISH staining experiment.

## Data availability

All data to support the findings of this study are included in the paper or supplementary information. The complete single-cell RNA-sequencing dataset can be accessed at https://betsholtzlab.org/Publications/LiverScRNAseq/search.html. The online resource will be made freely accessible upon publication. The raw sequencing data underlying the previously unpublished single-cell dataset will be deposited at *NCBI’s Gene Omnibus database (GEO)*, and made freely accessible upon publication, the accession number will be stated here. Previously published single-cell RNA-sequencing datasets used in this study are available at *GEO* under the accession numbers: GSE158183, GSE172492, GSE212047, GSE168933, GSE136103, and GSE137720. Further information about reagents and resources are available from the corresponding author, Lars Muhl upon reasonable request.

## Acknowledgements

We thank Professor Moustapha Hassan and the Pre-Clinical Laboratories (PKL)– Karolinska University Hospital Huddinge, as well as the Karolinska Institutet MedH fluorescent activated cell sorting facility, Cecilia Olsson, Pia Peterson, Veronica Sundell, Louise Larsson, Jana Chmielniakova, and Helene Leksell for technical help. We would also like to thank the Single-Cell Core Facility Flemingsberg campus, SICOF, which is supported by the infrastructure board of the Karolinska Institutet, for their single-cell sequencing services, and the Molecular Imaging Center (MIC), Department of Biomedicine, University of Bergen. We thank Dr. Lucile Miquerol for providing tissues from *Gja5^GFP^* reporter mice. This study was supported by grants from Magn. Bergvalls Foundation (L.M.: 2020-03735, 2021-04275, 2022-158), the Swedish Cancer Society (CB.: 21 1714Pj), the Swedish Research Council (C.B.: 2015-00550, K.G.: 2021-04896, M.A.M.: 2023-02655, U.L.: 2024-02414), Theme-based Research Scheme Hong Kong (U.L.: T12-712/21R), the K. and A. Wallenberg Foundation (C.B. and K.G.: 2020.0057), and the Leduq Foundation (C.B. and M.A.M.: 23CVD02; C.B.: 22CVD01).

## Supplementary Information

**Supplementary Figure 1 (related to Figure 1).**
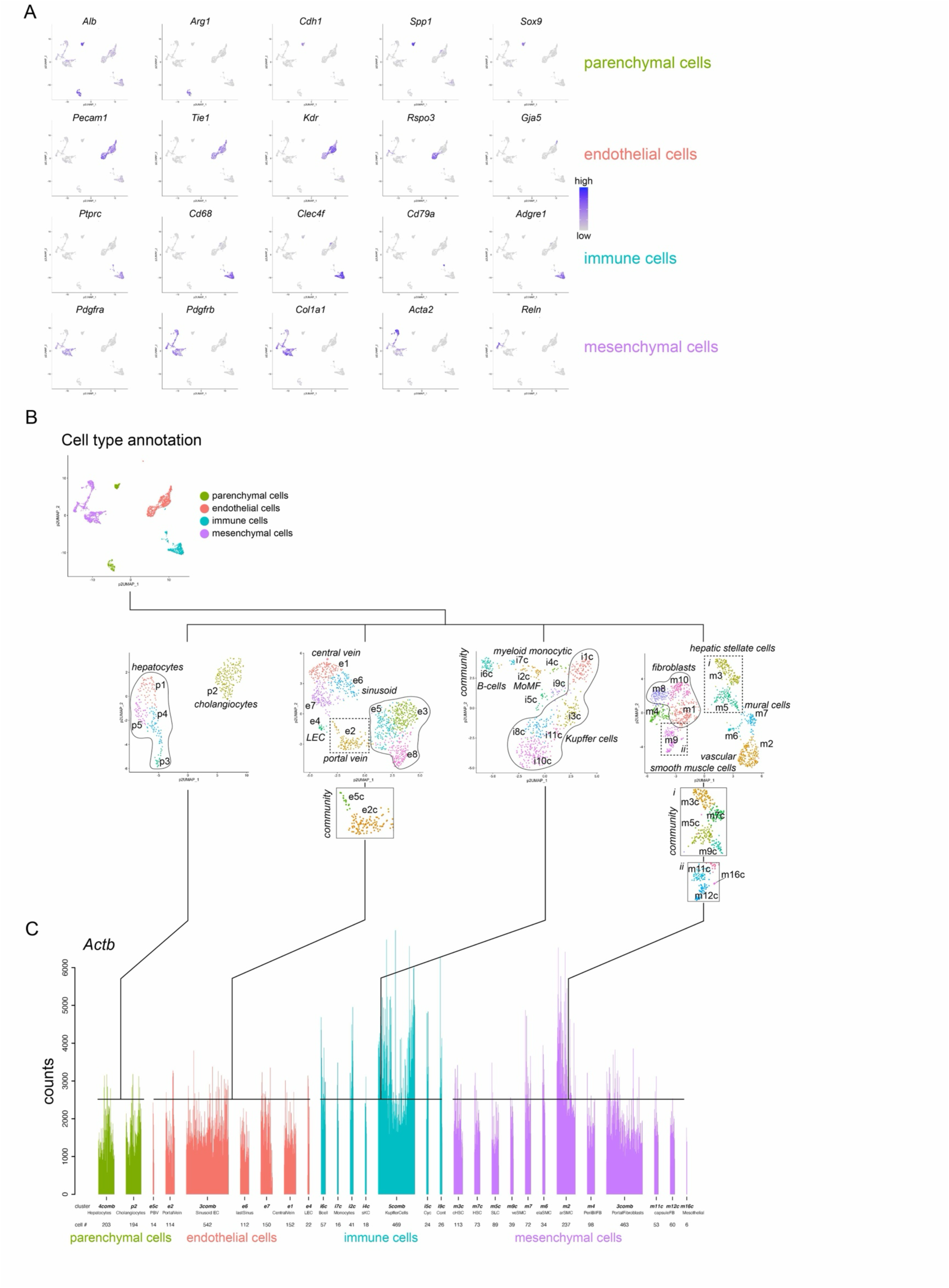
**A** UMAP visualization of the expression level of selected canonical marker genes used for cell type class identification. **B** UMAP visualization of the complete dataset color coded for cell type classes (upper panel) and UMAP visualizations of the separately analyzed cell class datasets, color coded for the clustering results using the multilevel or community setting as indicated. Cell type annotations are shown. Note that clusters #p1, p3, p4, and p5 in the parenchymal dataset, clusters #e3, e5, and e8 in the endothelial cell dataset, clusters #i1c, i3c, i8c, i10c, and i11c in the immune cell dataset, and clusters #m1, m8, and m10 in the mesenchymal cell dataset were combined before SPIN cell ordering and bar plot construction. **C** Bar plot visualization of the complete dataset using clustering and annotations as determined in the separate analyses of the cell class datasets. The bars are color coded for cell type classes. Note that the in-cluster cell order was calculated using the SPIN algorithm. The expression level of *Actb* is shown, each bar represents one single-cell transcriptome. The bar plot and UMAP visualization of the expression levels for each gene can be accessed at http://betsholtzlab.org/Publications/LiverScRNAseq/search.html.

**Supplementary Figure 2 (related to Figure 1 and 2).**
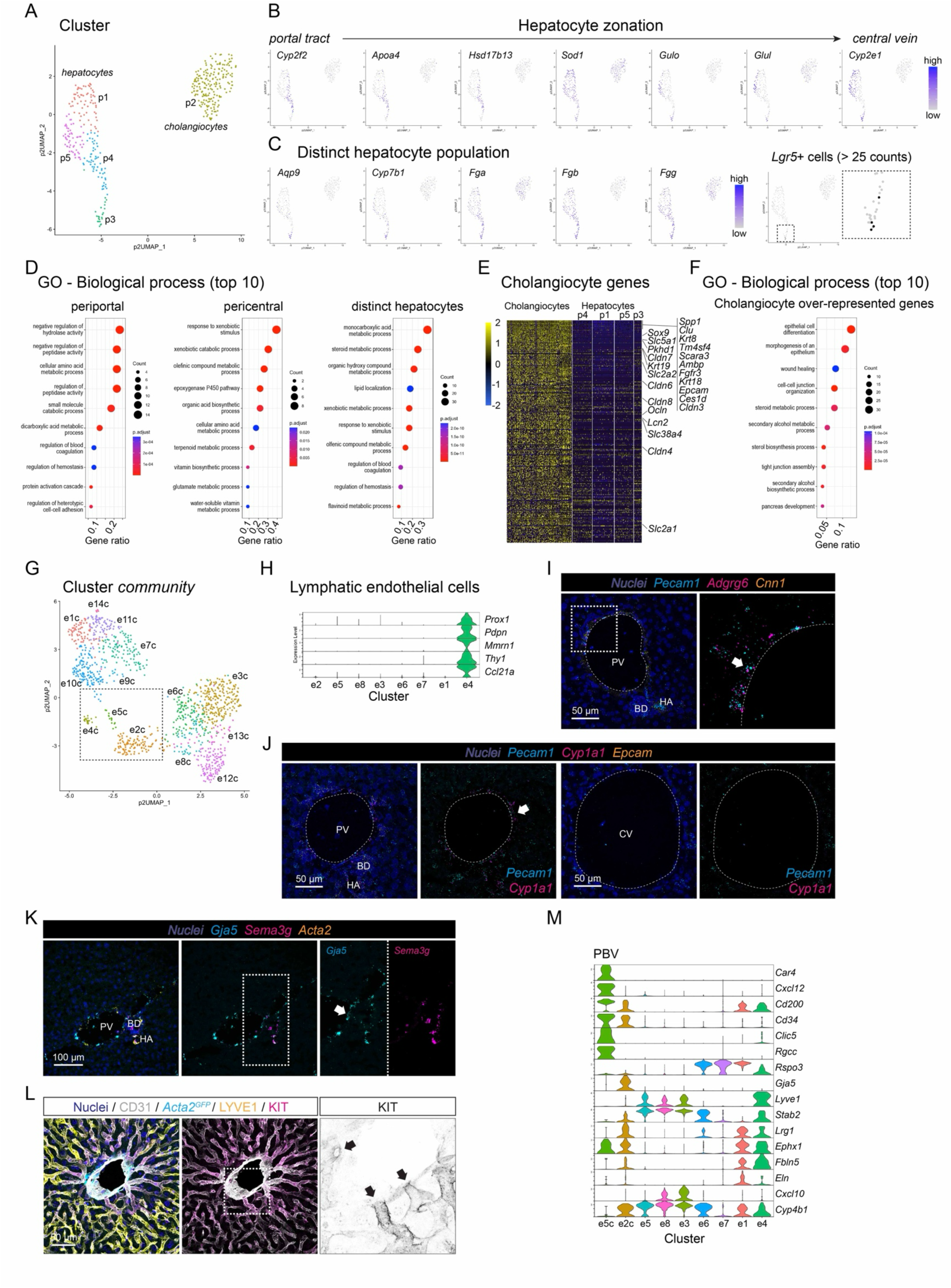
**A** UMAP visualization of the pagoda2 clustering result for the parenchymal cell dataset. **B** UMAP visualization of the expression level of selected genes identified as differentially expressed in hepatocytes along the portal-central axis. **C** UMAP visualization of exemplary genes with enriched expression in cluster #p3 and the highlighting of cells that express *Lgr5* (>25 counts). **D** Dot plots showing the top10 overrepresented GO terms of the genes enriched in the different hepatocyte subsets (periportal, left panel; pericentral, middle panel; distinct, right panel). **E** Heat map showing genes with enriched expression in cholangiocytes compared to hepatocytes. **F** Dot plot showing the top10 overrepresented GO terms of cholangiocyte enriched genes. **G** UMAP visualization of the clustering result of the EC dataset using the pagoda2 community setting. **H** Violin plot showing genes expressed in lymphatic endothelial cells (cluster #e4). **I** ISH for *Pecam1*, *Adgrg6*, and *Cnn1* on a liver tissue section. The arrow highlights the expression of *Adgrg6* in portal vein endothelial cells. **J** ISH for *Pecam1*, *Cyp1a1*, and *Epcam* on a liver tissue section. The arrow highlights the expression of *Cyp1a1* in portal vein endothelial cells. **K** ISH for *Gja5*, *Sema3g*, and *Acta2* on a liver tissue section. The arrow highlights the expression of *Gja5* in portal vein endothelial cells. **L** IF for CD31, LYVE1, and KIT on a liver tissue section from an *Acta2^GFP^* reporter mouse. The arrows highlight the terminal sinusoidal endothelial cells before entering the central vein. **M** Violin plot showing genes expressed by endothelial cells of the peribiliary vasculature (PBV, cluster #e5c) together with marker genes for large vessels (central vein, portal veins), as well as sinusoids. PV: portal vein, CV: central vein, HA: hepatic artery, BD: bile duct. Scale bars are indicated in the respective image panels.

**Supplementary Figure 3 (related to Figure 3).**
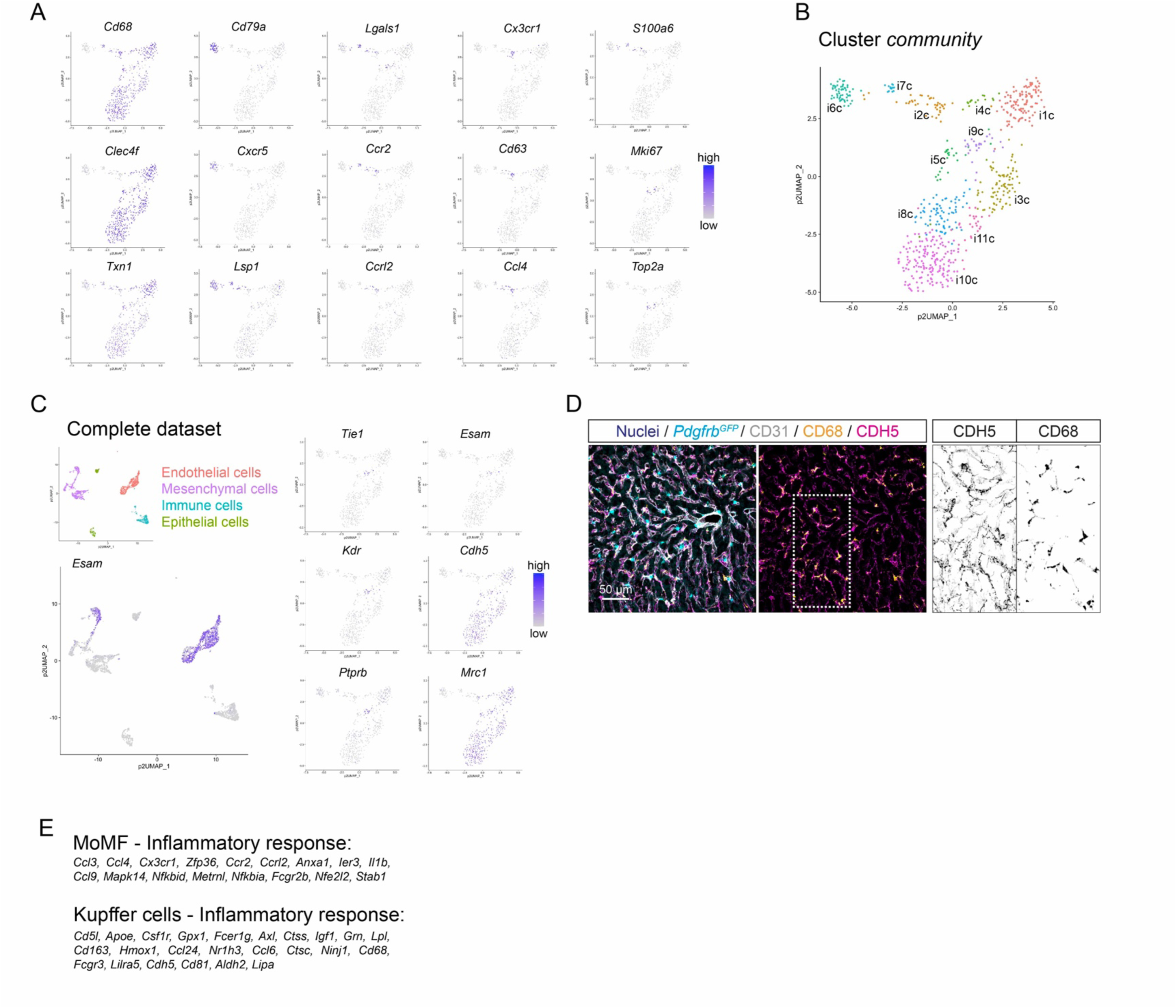
**A** UMAP visualization of the expression levels of exemplary genes with cluster-enriched expression in the immune cell subset. **B** UMAP visualization of the clustering result of the immune cell dataset using the pagoda2 community setting. **C** UMAP visualization of cell type distribution of the cell type classes in the complete dataset (upper left panel), the expression level of *Esam* in the complete dataset (left lower panel), and the expression levels of *Tie1*, *Esam*, *Kdr*, *Cdh5*, *Ptprb*, and *Mrc1* in the immune cell subset (right panel). Note in the visualization of the complete dataset that the expression of *Esam* is present in the endothelial cells and parts of the mesenchymal cells, but not in the immune cells. Only few transcriptomes in the immune cell subset show expression of *Esam*, which are also positive for the expression of other endothelial cell-specific transcripts, such as *Tie1*, *Kdr*, and *Ptprb*. **D** IF for CD31, CD68, and CDH5 on a liver tissue section from a *Pdgfrb^GFP^* reporter mouse. **E** Lists of genes belonging to the GO term ‘inflammatory response’ with enriched expression in either MoMFs (left panel) or Kupffer cells (right panel). Scale bars are indicated in the respective image panels.

**Supplementary Figure 4 (related to Figure 4).**
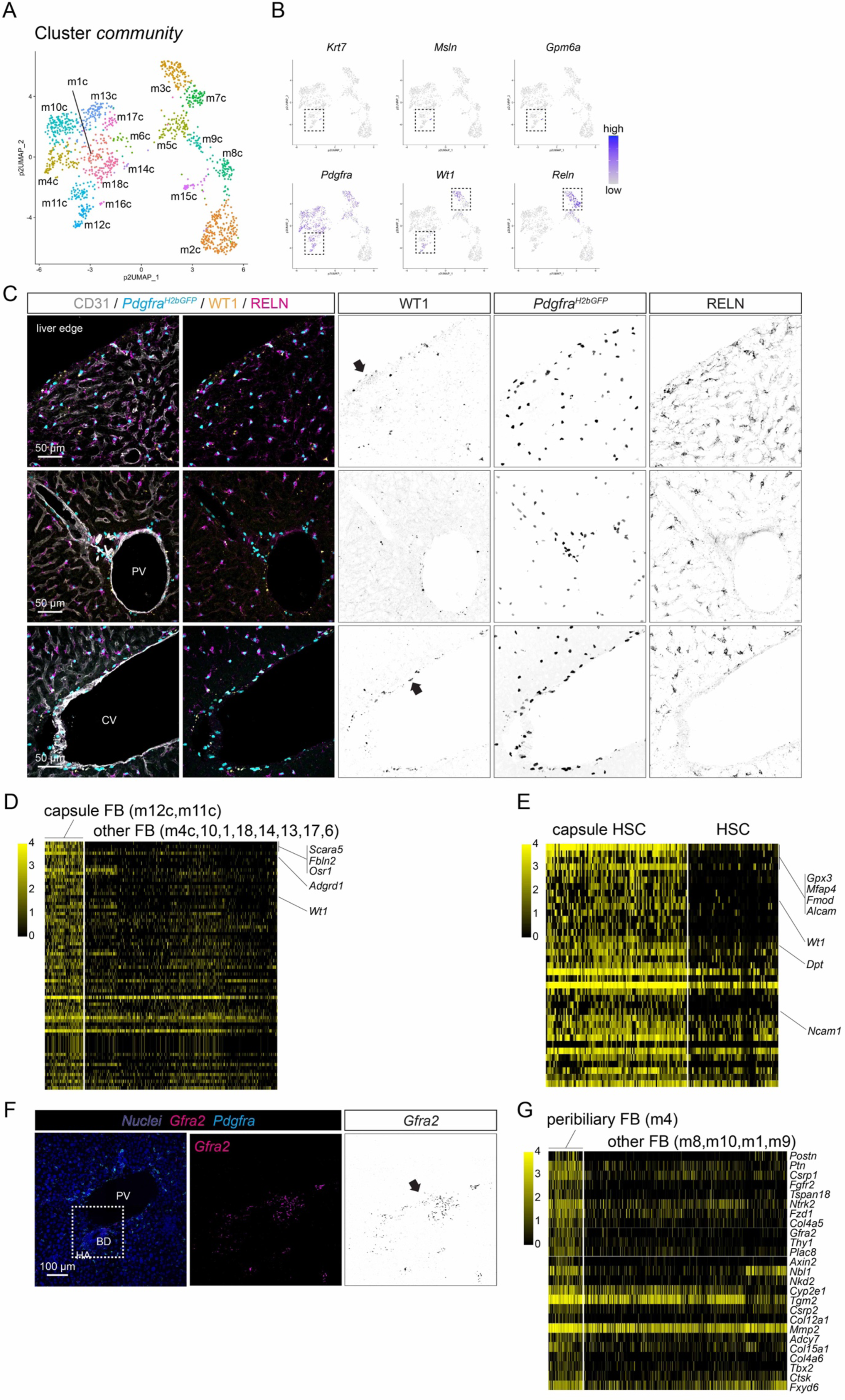
**A** UMAP visualization of the pagoda2 clustering result of the mesenchymal cell subset using the community setting. **B** UMAP visualization of the expression levels of selected genes with the area of pagoda2 multilevel clusters #m3 and m9 indicated. **C** IF for CD31, WT1, and RELN on a liver tissue sections from a *Pdgfra^H2bGFP^* reporter mouse focusing on the liver edge (upper panel), the portal tract (middle panel), or the central vein (lower panel). The arrows highlight WT1 positive cells. **D** Heat map showing the expression of genes that exhibit enriched expression in capsular fibroblasts (clusters #m11c, m12c), compared to all other fibroblast populations. **E** Heat map showing the genes that show enriched expression in cHSC (cluster #m3c), compared to HSC (cluster #m7c). **F** ISH for *Gfra2* and *Pdgfra* on a liver tissue section. The arrow indicates peribiliary fibroblasts. **G** Heat map showing the genes that are enriched expressed in peribiliary fibroblasts (cluster #m4), compared to all other fibroblast populations. PV: portal vein, BD: bile duct, HA: hepatic artery, CV: central vein. Scale bars are indicated in the respective image panels.

**Supplementary Figure 5 (related to Figure 5).**
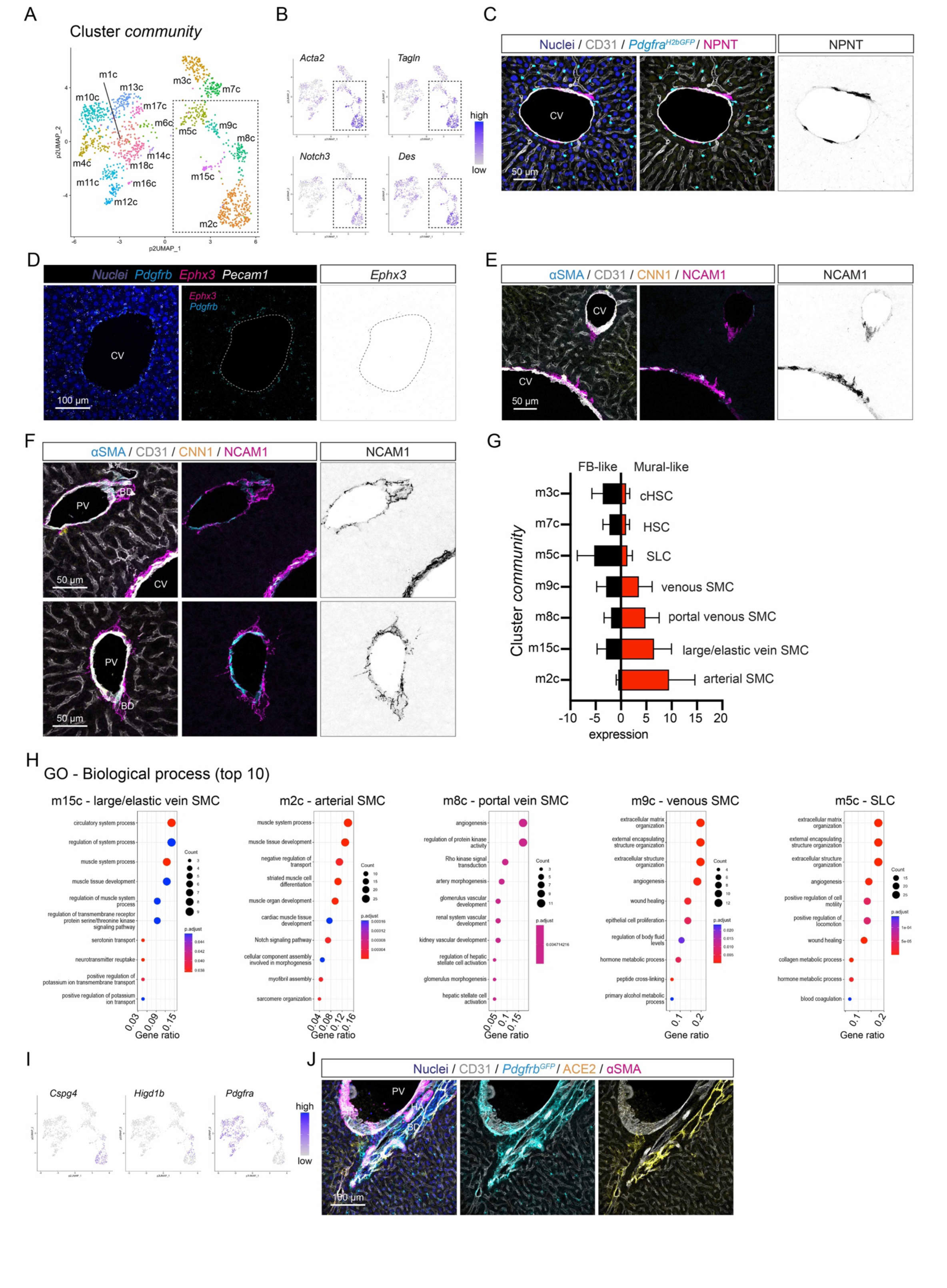
**A** UMAP visualization of the pagoda2 clustering result of the mesenchymal cell subset using the community setting with vascular mural cell clusters indicated. **B** UMAP visualization of the expression level of vascular SMC marker genes with vascular mural cell clusters indicated. **C** IF for CD31 and NPNT on a liver tissue section from a *Pdgfra^H2bGFP^* reporter mouse. **D** ISH for *Pdgfrb*, *Ephx3*, and *Pecam1* on a liver tissue section focusing on the central vein. **E** IF for αSMA, LYVE1, CNN1, and NCAM1 on a liver tissue section. **F** IF for αSMA, CD31, CNN1 and NCAM1 on a liver tissue section. **G** Calculation of fibroblast (FB)- or Mural cell-score from the 90-gene list ^19^ for community clusters containing HSC and vascular mural cell types. **H** Dot plot showing the top10 overrepresented GO terms for genes enriched in (from left to right) large-caliber vein SMC, arterial SMC, portal vein SMC, venous SMC, and SLC. **I** UMAP visualization of the expression level of pericyte marker genes (*Cspg4*, *Higd1b*, and *Pdgfrb*). **J** IF for CD31, ACE2, and αSMA on a liver tissue section from a *Pdgfrb^GFP^* reporter mouse focusing on the portal tract and bile duct. PV: portal vein, BD: bile duct, HA: hepatic artery, CV: central vein. Scale bars are indicated in the respective image panels.

**Supplementary Figure 6 (related to Figure 6).**
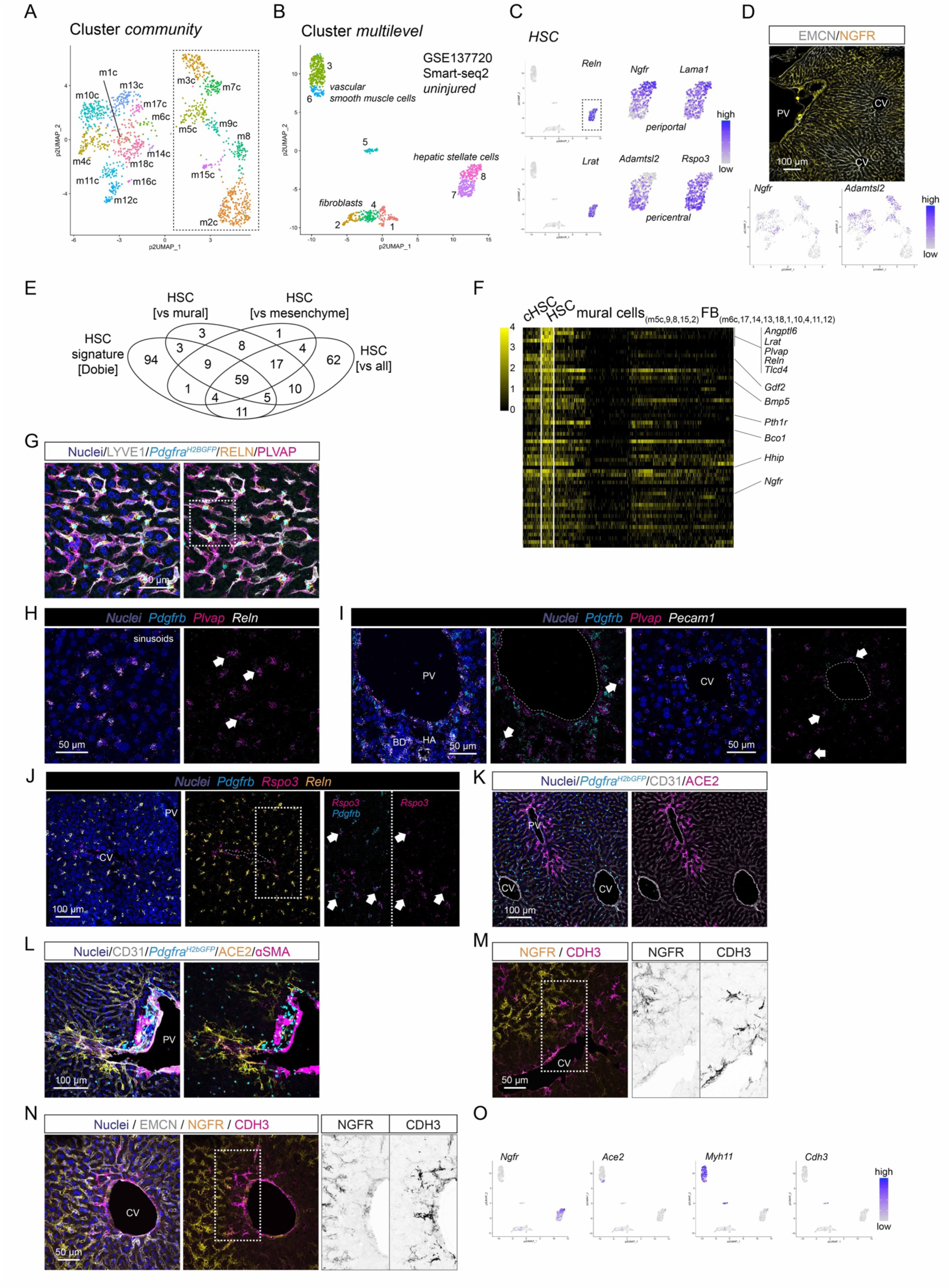
**A** UMAP visualization of the pagoda2 clustering result of the mesenchymal cell subset using the community setting with HSC and vascular mural cell clusters indicated. **B** UMAP clustering result of the Smart-seq2 dataset (GSE137720) from Dobie *et al*., using cells from uninjured mice (see original publication for details) and pagoda2 multilevel setting. **C** UMAP visualization of the expression level of HSC marker genes (*Reln* and *Lrat*), and HSC zonation markers (*Ngfr*, *Lama1*, *Adamtsl2*, and *Rspo3*) magnified in the indicated area of the Dobie *et al*. uninjured dataset. **D** IF for EMCN and NGFR on a liver tissue section (upper panel) and UMAP visualization of the expression level of *Ngfr* and *Adamtsl2* in the mesenchymal cell subset (lower panel). **E** Venn diagram showing the overlap of HSC-enriched genes computed from the Dobie *et al*. uninjured dataset, the mesenchymal cell subset (vs. mural, vs. mesenchyme), and the complete dataset (vs. all) (see also Table 1). Note that 59 genes are commonly detected as enriched expressed in HSC (cluster #m7c) in all four comparisons. **F** Heat map showing the expression level of the 59 HSC enriched genes in the mesenchymal cell subset. **G** IF for LYVE1, RELN, and PLVAP on a liver tissue section from a *Pdgfra^H2bGFP^* reporter mouse. The indicated frame is shown magnified in the Figure 6F. **H** ISH for *Pdgfrb*, *Plvap*, and *Reln* on a liver tissue section focusing on the sinusoids. Arrows indicate *Pdgfrb Plvap* double positive HSC. **I** ISH for *Pdgfrb*, *Plvap*, and *Pecam1* on liver tissue section focusing on the portal tract (left panels) or central vein (right panels). Arrows indicate *Pdgfrb Plvap* double positive HSC. **J** ISH for *Pdgfrb*, *Rspo3*, and *Reln* on a liver tissue section. Arrows indicate *Pdgfrb Rspo3* double positive HSC. **K** IF for CD31 and ACE2 on a liver tissue section from a *Pdgfra^H2bGFP^* reporter mouse. **L** IF for CD31, ACE2, and αSMA on a liver tissue section from a *Pdgfra^H2bGFP^* reporter mouse focusing on the portal tract. **M** IF for NGFR and CDH3 on a liver tissue section focusing on the central vein. **N** IF for EMCN, NGFR, and CDH3 on a liver tissue section focusing on the central vein. Note that NGFR and CDH3 signals are rarely overlapping. **O** UMAP visualization of the expression level of HSC subset markers (*Ngfr*, *Ace2*, *Myh11*, and *Cdh3*) in the Dobie *et al*. uninjured dataset. PV: portal vein, BD: bile duct, HA: hepatic artery, CV: central vein. Scale bars are indicated in the respective image panels.

**Supplementary Figure 7 (related to Figure 7).**
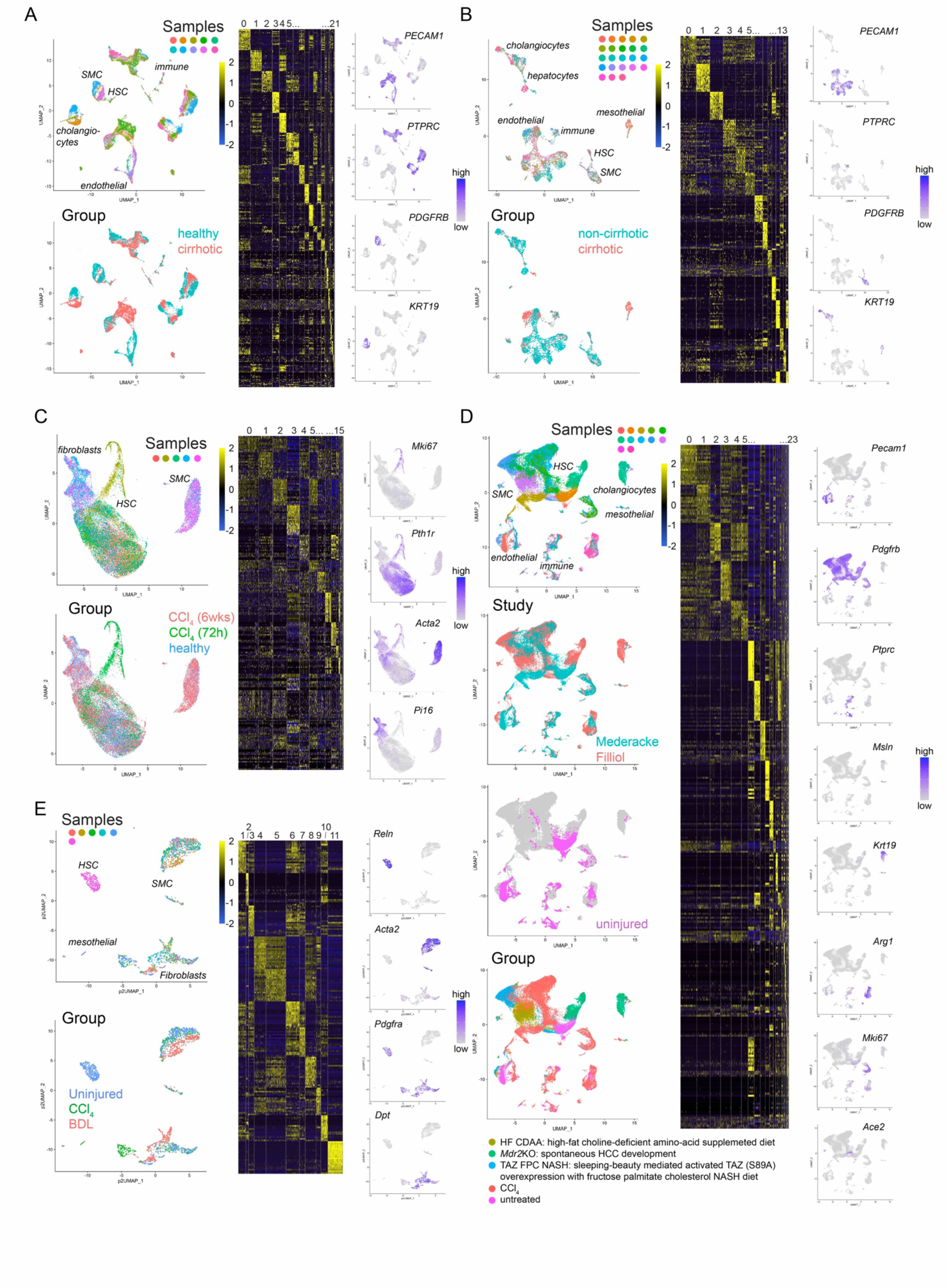
**A** UMAP visualization showing the distribution of the different samples (upper left panel), or the distribution of cells from the healthy or cirrhotic sample groups (lower left panel), and heat map showing the top20 enriched genes for each cluster (middle panel), as well as UMAP visualization showing the expression level of cell type class marker genes (right panel) in the Ramachandran *et al*. human 10X dataset. **B** UMAP visualization showing the distribution of the different samples (upper left panel), or the distribution of cells from the non-cirrhotic or cirrhotic sample groups (lower left panel), and heat map showing the top20 enriched genes for each cluster (middle panel), as well as UMAP visualization showing the expression level of cell type class marker genes (right panel) in the Bunomo *et al*. human 10X dataset. **C** UMAP visualization showing the distribution of the different samples (upper left panel), or the distribution of cells from the uninjured or CCl_4_ treated sample groups (lower left panel), and heat map showing the top20 enriched genes for each cluster (middle panel), as well as UMAP visualization showing the expression level of cell type marker genes (right panel) in the Dobie *et al*. mouse 10X dataset. **D** UMAP visualization showing the distribution of the different samples, the distribution of cells from the Mederacke *et al*., or Filliol *et al*. studies, and the distribution of cells from untreated samples, or the distribution of cells from all the different treatment groups (left panel from top to bottom), as well as heat map showing the top20 enriched genes for each cluster (middle panel), as well as UMAP visualization showing the expression level of cell type class marker genes (right panel) in the combined Mederacke *et al*. and Filliol *et al*. mouse 10X dataset. **E** UMAP visualization showing the distribution of the different samples (upper left panel), or the distribution of cells from the uninjured, CCl_4_ treated, or BDL treated sample groups (lower left panel), and heat map showing the top20 enriched genes for each cluster (middle panel), as well as UMAP visualization showing the expression level of mesenchymal cell type marker genes (right panel) in the complete Dobie *et al*. mouse Smart-seq2 dataset.

**Supplementary Table 1:** Antibodies and in situ probes.

**Primary antibodies**
| Protein | Gene | Species | Company | Reference | Comment |
| --- | --- | --- | --- | --- | --- |
| ACE2 | *Ace2* | goat | R&D Systems | AF3437 |  |
| alpha-SMA | *Acta2* | mouse | Sigma | C6198 | Cy3 conjugated |
| alpha-SMA | *Acta2* | mouse | Sigma | F3777 | FITC conjugated |
| CD200 | *Cd200* | goat | R&D Systems | AF2724 |  |
| CD31 | *Pecam1* | rat | BD Bioscience | 550274 |  |
| CD31 | *Pecam1* | goat | R&D Systems | AF3628 |  |
| CD31 | *Pecam1* | rat | BD Bioscience | 561814 | APC conjugated for FACS |
| CD31 | *Pecam1* | rat | BD Bioscience | 561813 | FITC conjugated for FACS |
| CD36 | *Cd36* | goat | R&D Systems | AF2519 |  |
| CD68 | *Cd68* | rabbit | Cell Signaling | 97778T |  |
| CD68 | *Cd68* | rat | BioLegend | 137008 | APC conjugated for FACS |
| CLEC4F | *Clec4f* | goat | R&D Systems | AF2784 |  |
| CLEC4F | *Clec4f* | mouse | BioLegend | 156803 | AlexaFluor-647 conjugated |
| CNN1 | *Cnn1* | rabbit | Abcam | ab216651 |  |
| EMCN | *Emcn* | rat | eBiosciences | 14-5851-82 |  |
| EPCAM (CD326) | *Epcam* | rat | eBiosciences | 17-5791-82 | APC conjugated for FACS |
| GPM6A | *Gpm6a* | rat | LS-Bio | LS-C179305 |  |
| KIT | *Kit* | goat | R&D Systems | AF1356 |  |
| LYVE1 | *Lyve1* | rabbit | Cell Signaling | 67538S |  |
| NCAM1 | *Ncam1* | goat | R&D Systems | AF2408 |  |
| NGFR | *Ngfr* | rabbit | Abcam | ab52987 |  |
| NPNT | *Npnt* | goat | R&D Systems | AF4298 |  |
| P-Cadherin | *Cdh3* | goat | R&D Systems | AF761 |  |
| PLVAP | *Plvap* | rat | BD Bioscience | 550563 |  |
| RELN | *Reln* | goat | R&D Systems | AF3820 |  |
| VE-Cadherin | *Cdh5* | goat | R&D Systems | AF1002 |  |
| VE-Cadherin | *Cdh5* | rat | eBiosciences | 53-1441-82 | AlexaFluor-488 conjugated |
| VE-Cadherin | *Cdh5* | rat | eBiosciences | 12-1441-82 | PE conjugated for FACS |
| VE-Cadherin | *Cdh5* | rat | eBiosciences | 48-1441-80 | eFluor450 conjugated for FACS |
| vWF | *Vwf* | rabbit | DAKO | A0082 |  |
| WT1 | *Wt1* | rabbit | Abcam | ab89901 |  |

**RNAscope probes**
| Gene | Species | Channel | Company | Reference |
| --- | --- | --- | --- | --- |
| *Ace2* | mouse (Mm) | C2 | ACD | 417081-C2 |
| *Acta2* | mouse (Mm) | C3 | ACD | 319531-C3 |
| *Adgrg6* | mouse (Mm) | C2 | ACD | 467351-C2 |
| *Ccn3 (Nov)* | mouse (Mm) | C1 | ACD | 415341 |
| *Chrdl1* | mouse (Mm) | C1 | ACD | 442811 |
| *Cnn1* | mouse (Mm) | C1 | ACD | 483791 |
| *Cyp1a1* | mouse (Mm) | C1 | ACD | 464611 |
| *Epcam* | mouse (Mm) | C2 | ACD | 418151-C2 |
| *Ephx3* | mouse (Mm) | C1 | ACD | 412511 |
| *Hhip* | mouse (Mm) | C2 | ACD | 448441-C2 |
| *Gfra2* | mouse (Mm) | C1 | ACD | 441481 |
| *Gja5* | mouse (Mm) | C2 | ACD | 518041-C2 |
| *Pdgfra* | mouse (Mm) | C2 | ACD | 480661-C2 |
| *Pdgfrb* | mouse (Mm) | C2 | ACD | 411381-C2 |
| *Pecam1* | mouse (Mm) | C1 or C3 | ACD | 316721, 316721-C3 |
| *Pecam1 O1* | mouse (Mm) | C2 | ACD | 471481-C2 |
| *Pln* | mouse (Mm) | C1 | ACD | 506241 |
| *Plvap* | mouse (Mm) | C1 | ACD | 440221 |
| *Reln* | mouse (Mm) | C3 | ACD | 405981-C3 |
| *Rspo3* | mouse (Mm) | C1 | ACD | 402011 |
| *Sema3g* | mouse (Mm) | C1 | ACD | 494691 |
| *Stab2* | mouse (Mm) | C1 | ACD | 406611 |
| *Wif1* | mouse (Mm) | C1 | ACD | 412361 |
| *Wt1* | mouse (Mm) | C1 | ACD | 432711 |

